# A heme-binding protein produced by *Haemophilus haemolyticus* inhibits non-typeable *Haemophilus influenzae*

**DOI:** 10.1101/626416

**Authors:** Roger D. Latham, Mario Torrado, Brianna Atto, James L. Walshe, Richard Wilson, J. Mitchell Guss, Joel P. Mackay, Stephen Tristram, David A. Gell

## Abstract

Many commensal bacteria and opportunistic pathogens scavenge heme from their environment. Pathogens and host are engaged in an arms race to control access to heme, but similar conflicts between bacterial species that might regulate pathogen colonisation are largely unknown. We show here that a commensal bacterium, *Haemophilus haemolyticus*, makes hemophilin, a heme-binding protein that not only allows the bacterium to effectively scavenge heme for its own growth, but also inhibits co-culture of the opportunistic pathogen, non-typeable *Haemophilus influenzae* (NTHi), by heme starvation. Knockout of the hemophilin gene abrogates the ability of *H. haemolyticus* to inhibit NTHi and an x-ray crystal structure shows that hemophilin has a previously unreported heme-binding structure. The bound heme molecule is deeply buried and the heme iron atom is coordinated through a single histidine side chain. Biochemical characterization shows that this arrangement allows heme to be captured in the ferrous or ferric state, and with small ferrous or ferric heme-ligands bound, suggesting hemophilin could function over in a wide range of physiological conditions. Our data raise the possibility that competition for heme between commensal and pathogenic bacteria can influence bacterial colonisation, and therefore disease likelihood, and suggest that strains of *H. haemolyticus* that overproduce hemophilin might have therapeutic uses in reducing colonisation and subsequent opportunistic infection by NTHi.

## Introduction

Non-typeable *Haemophilus influenzae* (NTHi) are Gram-negative bacteria with their reservoir in the upper respiratory tract of humans. Although frequent colonizers of healthy children and adults (Lemon et al., 2010), NTHi have replaced *H. influenzae* capsular type b (Hib) as the leading cause of invasive infections for this species (Van Eldere et al., 2014), and are also an important cause of non-invasive disease, such as recurrent otitis media (Ngo et al., 2016) and exacerbations of chronic obstructive pulmonary disease (Sethi and Murphy, 2008).

The management of NTHi infections is becoming increasingly difficult. The intrinsic heterogeneity of NTHi has hampered vaccine development, and despite significant effort, an effective vaccine is not currently available (Cerquetti and Giufre, 2016). There has been an increase in both the incidence and spectrum of antimicrobial resistance in NTHi (Tristram et al., 2007) and the development of new antibiotics for NTHi has recently been listed as a priority by the World Health Organisation, emphasising the need for new approaches to the prevention and management.

As an alternative to antibiotics, bacteriocin-producing strains of upper respiratory tract commensal streptococci have been commercialized as probiotics to prevent and treat *S. pyogenes* infections (Di Pierro et al., 2016), and, more recently, bacteriocin-producing strains of *Staphylococcus lugdunensis* were shown to be associated with reduced nasal carriage of *S. aureus* (Zipperer et al., 2016). In light of these studies, we considered that certain commensal *Haemophilus* spp. might have potential as probiotics to counter NTHi infection. *Haemophilus haemolyticus* are bacteria that are resident in the upper respiratory tract of healthy adults and children at sites that are also colonised by NTHi. *H. haemolyticus* is genetically and phenotypically closely related to NTHi, with both requiring heme and NAD for growth. However, unlike NTHi, *H. haemolyticus* is not considered to be a pathogen (Murphy et al., 2007; Zhang et al., 2014).

Recently, with the purpose of developing strains of *H. haemolyticus* as respiratory tract probiotics, we performed a screen of 100 *H. haemolyticus* isolates and found that the isolate BW1 produced a compound capable of inhibiting the growth of NTHi *in vitro* (Latham et al., 2017). Here, we show that the NTHi inhibitory factor is a heme binding protein, which we have named hemophilin. An x-ray crystal structure shows that hemophilin has a previously undescribed heme-binding fold. Recombinant hemophilin, and a hemophilin gene knockout in *H. haemolyticus*, demonstrate that hemophilin is sufficient and necessary for NTHi inhibition *in vitro*. Together, the results presented below suggest that hemophilin has an important role in heme uptake by *H. haemolyticus*, and that growth-inhibition of competing NTHi occurs by heme starvation.

## Results

### Identification of the hemophilin gene from two *Haemophilus haemolyticus* clinical isolates

We previously showed that an NTHi-inhibitory protein was produced by *H. haemolyticus* isolate BW1 (Latham et al., 2017). Further screening identified a second *H. haemolyticus* isolate, RHH122, with similar activity (Supplementary Figs. 1 and 2). To identify the NTHi-inhibitory protein, which we now call hemophilin, conditioned growth medium from stationary phase cultures of BW1 and RHH122 was fractionated by ammonium sulfate precipitation, size exclusion chromatography (SEC) and reversed-phase HPLC (RP-HPLC). Identical (mock) separation steps were performed for two control *H. haemolyticus* isolates (BW39 and BWOCT3) that lacked NTHi-inhibitory activity. For BW1 and RHH122 samples, RP-HPLC peaks containing NTHi-inhibitory activity eluted at the same retention time, and these peaks were absent from BW29 and BWOCT3 samples (Fig. 1A). SDS-PAGE and silver staining revealed one predominant polypeptide of ~30 kDa in the active fractions from BW1 and RHH122 (Fig. 1B and Supplementary Fig. 2), which was identified as a hypothetical protein (GenBank accession EGT80255) from *H. haemolyticus* strain M19107 by mass spectrometry and peptide mass fingerprint analysis (Fig. 1C). Peptide mass fingerprint analysis of whole RP-HPLC fractions identified EGT80255 peptides as the most abundant ions in samples with NTHi-inhibitory activity obtained from BW1 and RHH122, whereas MS analysis of matched RP-HPLC fractions from control strains, BW29 and BWOCT3, did not identify EGT80255 peptides above background (normalised intensity < 1%; Supplementary Table 1). Notably, no peptides corresponding to the first 22 amino acids of EGT80255 were identified, suggesting that a predicted signal peptide (Fig. 1C, blue) had been cleaved to release the mature hemophilin protein into the *H. haemolyticus* growth medium.

**Fig. 1.**
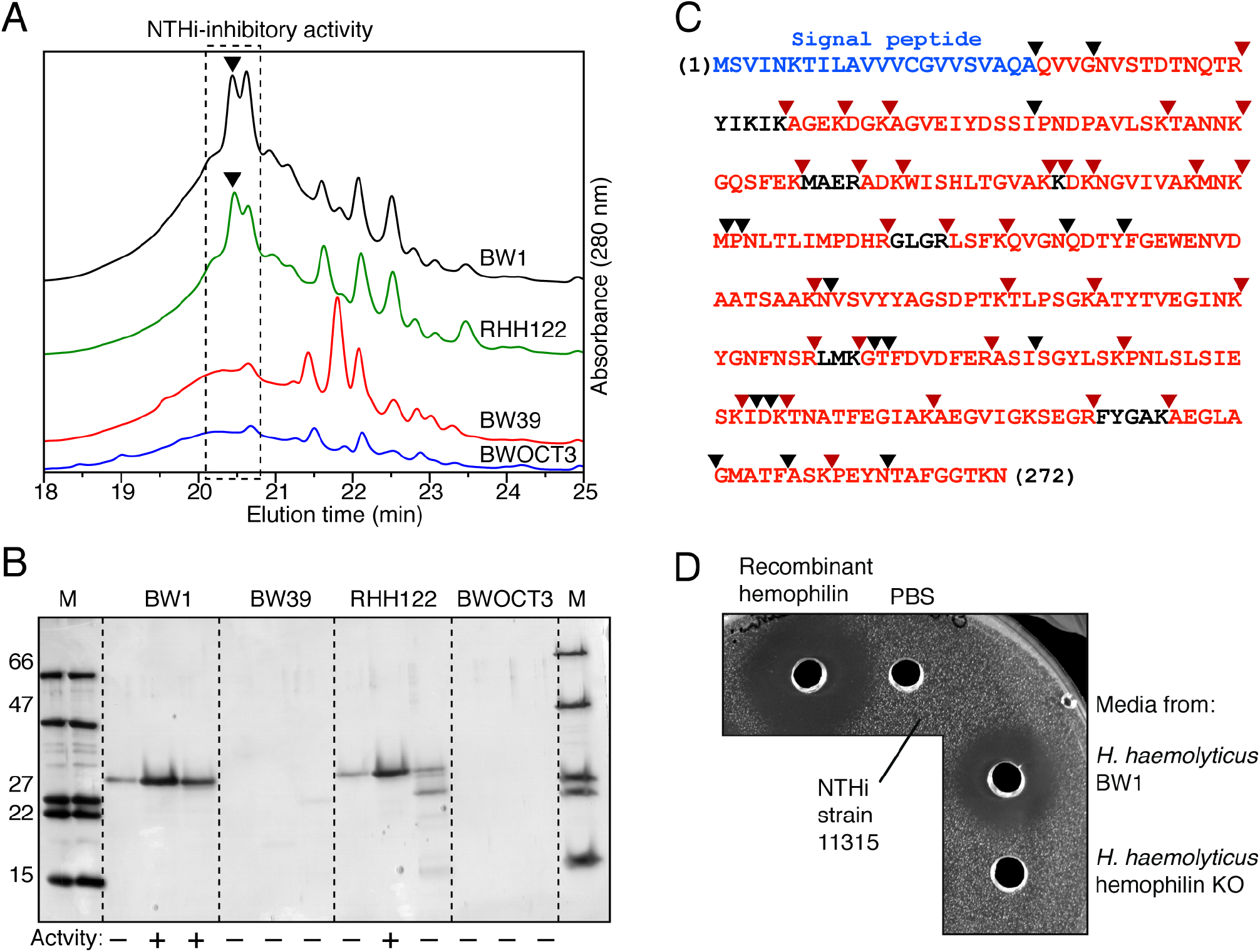
Hemophilin is a 27-kDa protein isolated from *H. haemolyticus* that inhibits the growth of NTHi. **(A)** RP-HPLC profiles show that NTHi-inhibitory activity (filled triangles) coincided with elution peaks that were present in BW1 and RHH122 samples and absent from control strains BW39 and BWOCT3. **(B)** Tris-tricine SDS-PAGE and silver staining of RP-HPLC fractions indicated by a dashed box in A. Markers (M; kDa); activity (+/−) from NTHi inhibition assay. **(C)** Tryptic peptides of hemophilin matching 92% of the sequence (red) for hypothetical protein EGT80255 were identified by MS; trypsin cleavage (red triangles) and other fragmentation sites (black triangles) are indicated. No peptides were matched to a putative signal peptide (blue). **(D)** Agar well diffusion assay. Activity abtained from: recombinant hemophilin (400 pmole; 20 μM); phosphate buffered saline (PBS); 1 mL of *H. haemolyticus* BW1 culture (OD_600_ = 0.88); 1 mL of *H. haemolyticus* hemophilin knockout mutant culture (OD_600_ = 1.1). The indicator strain is NTHi 11315.

**Fig. 2.**
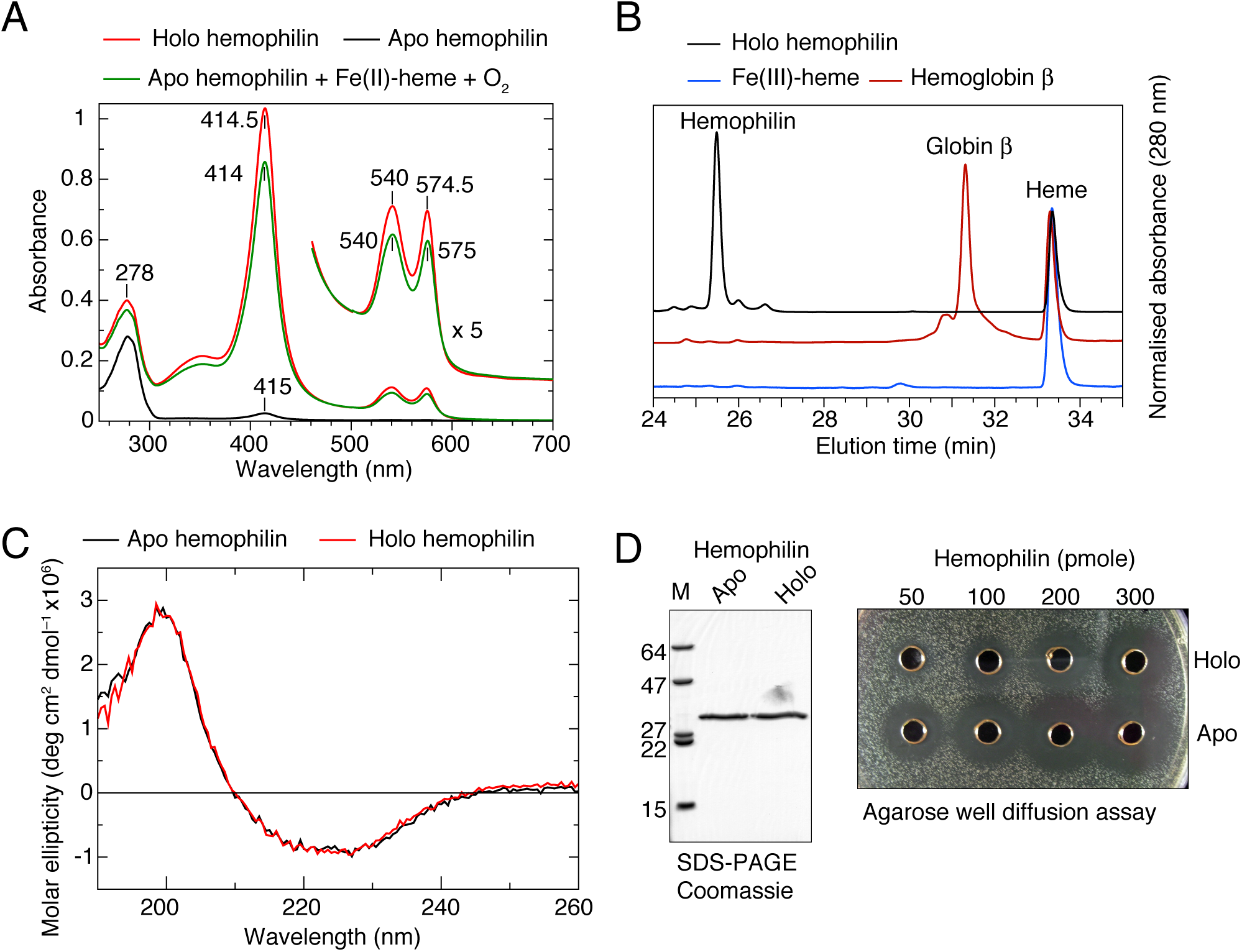
Hemophilin is a heme-binding protein. **(A)** UV-visible absorption spectra of recombinant holo (red) and apo hemophilin (black). Addition of ferrous heme and O_2_ to apo hemophilin yields a spectrum (green) essentially identical to that of holo hemophilin. **(B)** RP-HPLC shows heme is non-covalently bound to hemophilin; the retention time for the dissociated prosthetic group is identical to that of heme dissociated from human hemoglobin β chain, or purified heme. **(C)** CD spectra of holo and apo hemophilin (25 mM sodium phosphate, 125 mM NaF, pH 7.1) are highly similar, suggesting no large change in secondary structure upon heme binding. **(D)** Apo hemophilin has greater NTHi-inhibitory activity than the holo protein.

PCR-based sequencing of the hemophilin gene loci from BW1 and RHH122 genomic DNA confirmed the presence of ORFs that were identical to EGT80255. We cloned the hemophilin ORF from BW1 and expressed the mature form (residues 23−272, without the N-terminal signal peptide), in *E. coli*. Purified recombinant hemophilin displayed NTHi-inhibitory activity at micromolar concentrations (Fig. 1D). To show that hemophilin is the NTHi-inhibitory activity in *H. haemolyticus*, we generated a hemophilin gene knockout mutant of *H. haemolyticus* BW1 by insertional inactivation with a kanamycin resistance cassette. Media recovered from cultures of the knockout strain displayed no inhibitory activity against NTHi (Supplementary Fig. 3). Growth medium recovered from BW1 or the BW1 hemophilin gene knockout contained similar protein content overall, as judged by chromatographic separations and silver staining, with the notable exception that the hemophilin protein band was absent from the knockout samples (Supplementary Fig. 3). In summary, these results show that hemophilin is the NTHi-inhibitory protein from *H. haemolyticus* isolates BW1 and RHH122.

### Hemophilin is a heme binding protein

The UV-visible absorption spectrum of the hemophilin had strong absorption bands at 415 nm, 540 nm and 575 nm, which are characteristic of the Soret and α/β bands of a hemoprotein (Fig. 2A, red trace). The presence of non-covalently bound heme *b* was confirmed by RP-HPLC and mass spectrometry (Fig. 2B). Apo hemophilin (hemophilin with the heme group removed) was able to bind reduced ferrous heme or oxidised ferric heme, and the reconstituted holo proteins were competent to bind to a variety of small ligands that are specific for ferrous (O_2_, CO; Supplementary Fig. 4) or ferric (CN^−^, HS^−^; Supplementary Fig. 5) hemes. The UV-visible spectrum of apo hemophilin (Fig. 2A, black trace) reconstituted with ferrous heme and O_2_ (Fig. 2A, green) is essentially identical to that of hemophilin purified from *E. coli* as the holo protein (Fig. 2A, red), suggesting that the *E. coli* derived protein is the O_2_-bound form.

The holo and apo hemophilin fractions gave very similar far-UV CD spectra, with maxima at ~200 nm and minima between 220−230 nm, indicating a mixture of β-sheet and α-helical secondary structure elements, and no major change in secondary structure upon heme binding (Fig. 2C). This is unlike hemoglobins or heme-binding enzymes, which typically undergo at least partial denaturation in the absence of the heme cofactor, and is more similar to the apo state characteristics of transient heme binding proteins, such as heme transport proteins, which essentially have folded apo protein structures (Smith et al., 2010). In addition, the hemophilin apo protein displayed NTHi-inhibitory activity that was typically 3−4 fold higher than that of the holo protein (Fig. 2D). The structure and activity of apo hemophilin suggest that reversible binding to heme is likely to be a function of this protein.

### Hemophilin has a novel heme binding structure

Holo hemophilin was crystallized and the structure determined by x-ray diffraction to a resolution of 1.6 Å (Supplementary Table 2). Initial phases were obtained by single wavelength anomalous diffraction (SAD) from the heme iron, above the absorption K-edge. All residues of hemophilin (23−272) are visible in the electron density. The hemophilin structure comprises an N-terminal region with mixed α/β secondary structure (Fig. 3, blue–green) that binds a single heme molecule (Fig. 3A, grey sticks and orange sphere), and a C-terminal 8-stranded β-barrel (Fig. 3, yellow–red). Helix and loop insertions between the β-3 and β-4 strands, and between the β-5 and β-6 strands, cover both faces of the porphyrin ring and provide the majority of protein-heme contacts. The dali server shows that several bacterial proteins have structural similarity to hemophilin (Supplementary Fig. 6). Notably, all are cell-surface proteins with ligand binding functions (Supplementary Fig. 7), found in *Neisseriaceae* and some members of the *Pasteurellaceae*, but none have heme binding function or structural similarity within the hemophilin heme-binding site. No heme-binding domain with similarity to hemophilin could be detected by structure- or sequence-based searches.

**Fig. 3.**
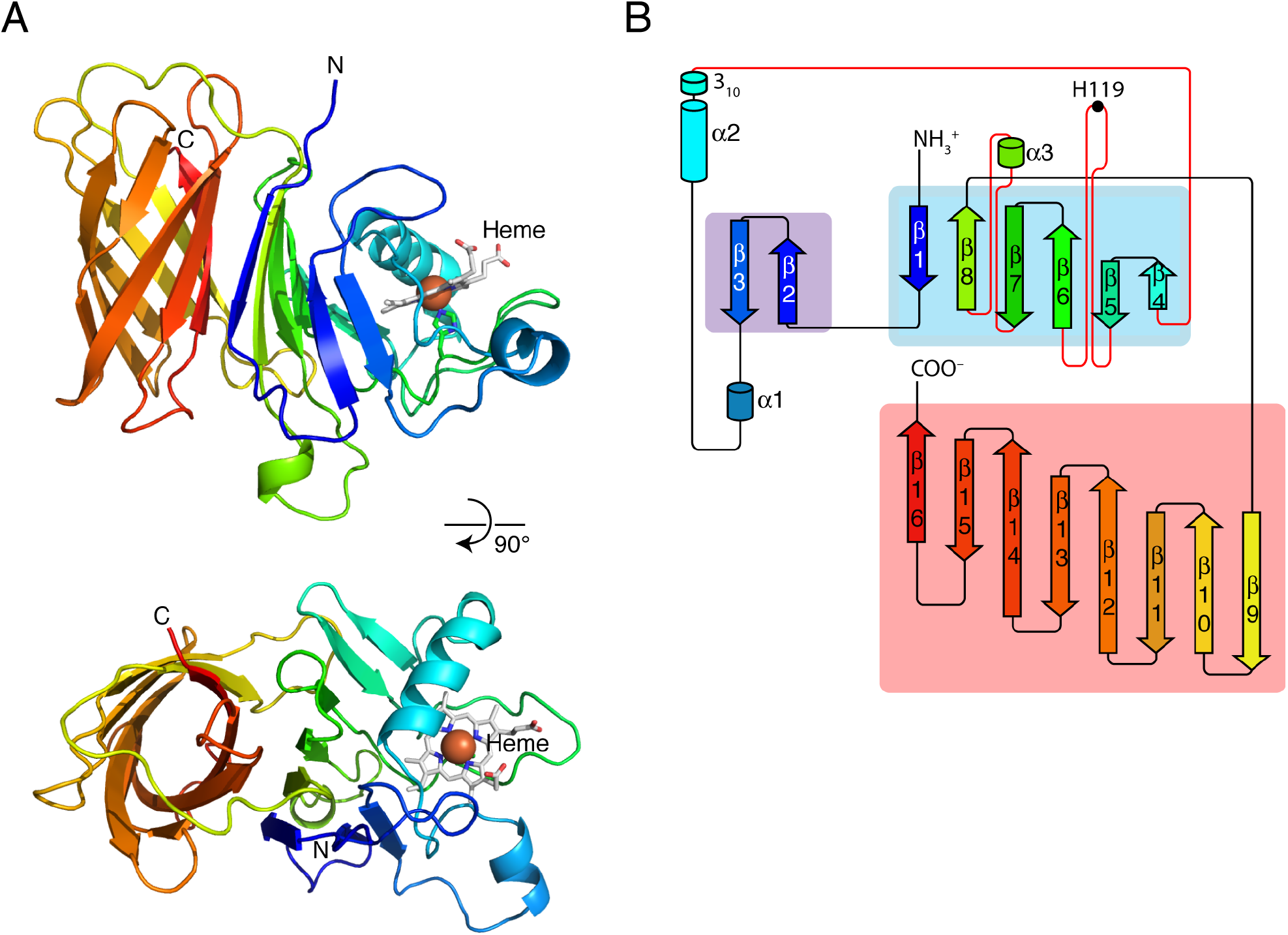
Hemophilin is a new heme-binding fold. **(A)** Richardson diagrams of the hemophilin crystal structure in two orthogonal views. **(B)** Topology diagram of hemophilin prepared based on the output from PROORIGAMI (Stivala et al., 2011).

In the hemophilin crystal structure, a heme molecule and a small exogenous heme ligand are clearly visible in electron difference density OMIT maps (Fig. 4A; electron difference density maps from the early stages of structure refinement using a model comprising only the hemophilin polypeptide are shown in Supplementary Fig. 8). Electron density and an anomalous diffraction peak at the ligand position indicated a single heavy atom, which was modelled as a chloride ion (Fig. 4A). The recombinant hemophilin protein used for crystallization was predominantly ferrous O_2_ complex but contained some ferric protein (Supplementary Fig. 9), and the crystallization conditions (pH 4.5) favoured further heme oxidation (Supplmentary Fig. 10), which would allow chloride binding (Supplementary Figure 11 shows that ferric hemophilin has a *K*_d_ of ~3 mM for chloride binding; chloride concentration in the crystallization drop was estimated to be 35–70 mM). The chloride refined at full occupancy and *B*-factors were consistent with those of the iron and other surrounding atoms. The Fe–Cl bond length in the hemophilin structure is 2.47(8) Å, which is in the range observed for Fe–Cl distances in heme protein structures (Kuwada et al., 2011; Singh et al., 2012; Kumar et al., 2013) (Supplementary Fig. 12; Supplementary Table 3).

**Fig. 4.**
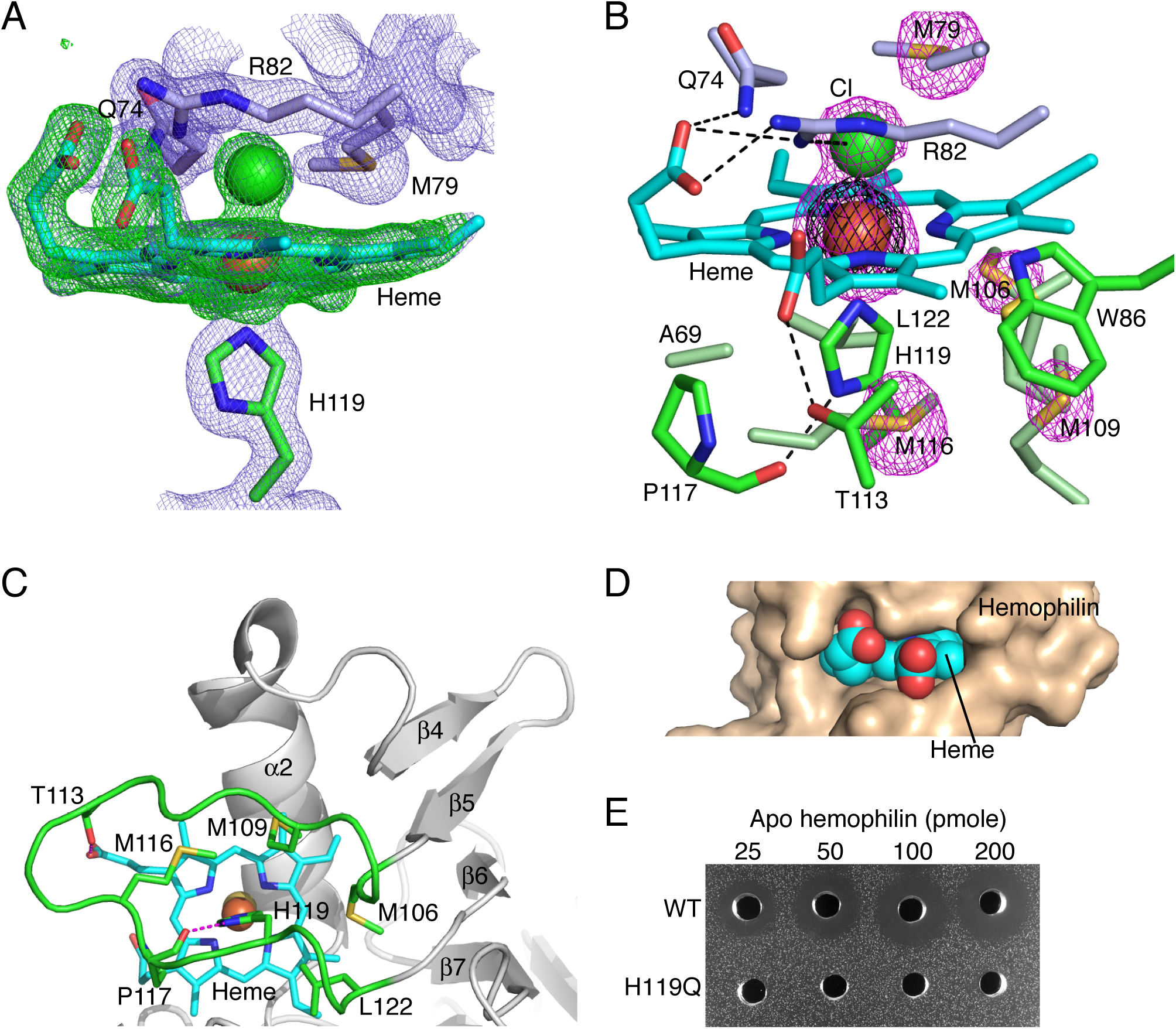
Features of the hemophilin heme pocket. **(A)** Figure shows *2F*_*O*_–*F*_*C*_ electron density map (contoured at 2 σ; blue mesh) and *F*_*O*_–*F*_*C*_ OMIT map (contoured at 4σ; refined without the heme and heme ligand; green mesh). **(B)** Protein side chains make extensive contacts with both faces of the porphyrin (cyan sticks). Anomalous diffraction map contoured at 25 σ (black mesh) identifies the position of the heme iron atom (orange sphere), and contoured at 5 σ (pink mesh) identifies sulfur atoms (yellow) of methionines, and a chloride ion (green sphere). **(C)** All the molecular contacts between hemophilin and the H119-proximal side of the heme are provided by a single protein loop (green). **(D)** The heme group (spheres) is buried with the central iron obscured. **(E)** Mutation of the heme-coordinating histidine (H119Q) leads to loss of NTHi-inhibtory activity.

The heme iron in hemophilin is coordinated in one axial position by the Nε2 atom of His119. The His119 side chain also makes a hydrogen bond through Nδ1 to the backbone carbonyl of Pro117 and is surrounded by a hydrophobic cage comprising Ala69, Met106, Met109, Met116, Pro117, Leu122 side chains (Fig. 4B) that shields one face of the porphyrin and the Fe–Nε bond from water, providing a similar physicochemical environment to that of the proximal His in metazoan hemoglobins. His119 and all its caging residues, bar Ala69, lie on the protein loop between β strands 5 and 6 that extends across one face of the heme group (Fig. 4C); a conformational change in this loop could, therefore, be a possible mechanism to achieve heme entry and exit—solution state studies will be required to address this. The heme Fe atom sits in the plane of the porphyrin— 0.04(8) Å from the least squares plane defined by the pyrrole N atoms—consistent with a 6-coordinate octahedral geometry. The side chain of Arg82 extends across the distal face of the porphyrin with the guanidinium group making cation-π and π-π interactions (Kumar et al., 2018) with the porphyrin, and hydrogen bonding interactions with the heme ligand and the porphyrin 17-propionate (Fig. 4B); a similar Arg conformation is seen in other heme protein structures with chloride ligands (Supplementary Fig. 12). The heme is oriented with the iron atom and porphyrin ring almost completely buried (solvent accessible non-polar area of 47 Å^2^, compared to 688 Å^2^ for free heme) and the ionisable propionate groups pointing out into solvent (Fig. 4D).

To investigate whether the heme binding function of hemophilin is required for NTHi-inhibitory activity, we made a His to Gln substitution of the heme-coordinating His119 (H119Q). Unlike wild-type hemophilin, the H119Q mutant protein purified from *E. coli* without a heme cofactor, indicating a large reduction in heme binding affinity (Supplementary Fig. 13). Mixing heme with the H119Q mutant gave a spectrum with peaks at 404, 485 and 602 nm, similar to spectra of ferric heme:protein complexes without an Fe-coordinating side chain (Gao et al., 2018), suggesting that the H119Q mutant binds heme, but probably does not form an axial bond to the iron. The H119Q mutant was not inhibitory to NTHi at the concentrations tested (Fig. 4E and Supplementary Fig. 13), indicating that heme coordination through H119 contributes substantially to NTHi inhibition.

### Hemophilin sequesters a pool of heme and makes it available to *H. haemolyticus* but not to the related species NTHi

*H. haemolyticus* is unable to synthesise porphyrin and must acquire heme from the environment (Norskov-Lauritsen, 2014). Therefore, we hypothesised that hemophilin might function as part of a heme uptake pathway. To investigate this idea, we compared the growth of BW1 and the hemophilin knockout strain under different heme supplementation regimes. BW1 that was propogated in heme-replete TSB media, continued to grow when heme supplementation was withdrawn (Fig. 5A), presumably due to accumulation of heme or porphyrin stores under prior heme-replete conditions, consistent with previous reports (Mason et al., 2011; Vogel et al., 2012). After a 14-hour conditioning period in heme-deficient TSB media, BW1 now showed a strong dependence on heme supplementation for growth (Fig. 5B and Supplementary Fig. 14). In contrast, after conditioning in heme-replete TSB, the hemophilin knockout strain showed poor growth at all heme concentrations (Fig. 5C), and this was exacerbated after heme starvation (Fig. 5D), whereas the strain could be propogated for long periods on blood-supplemented media (not shown). These results suggest that the hemophilin knockout strain has a reduced capacity to utilise free heme.

**Fig. 5.**
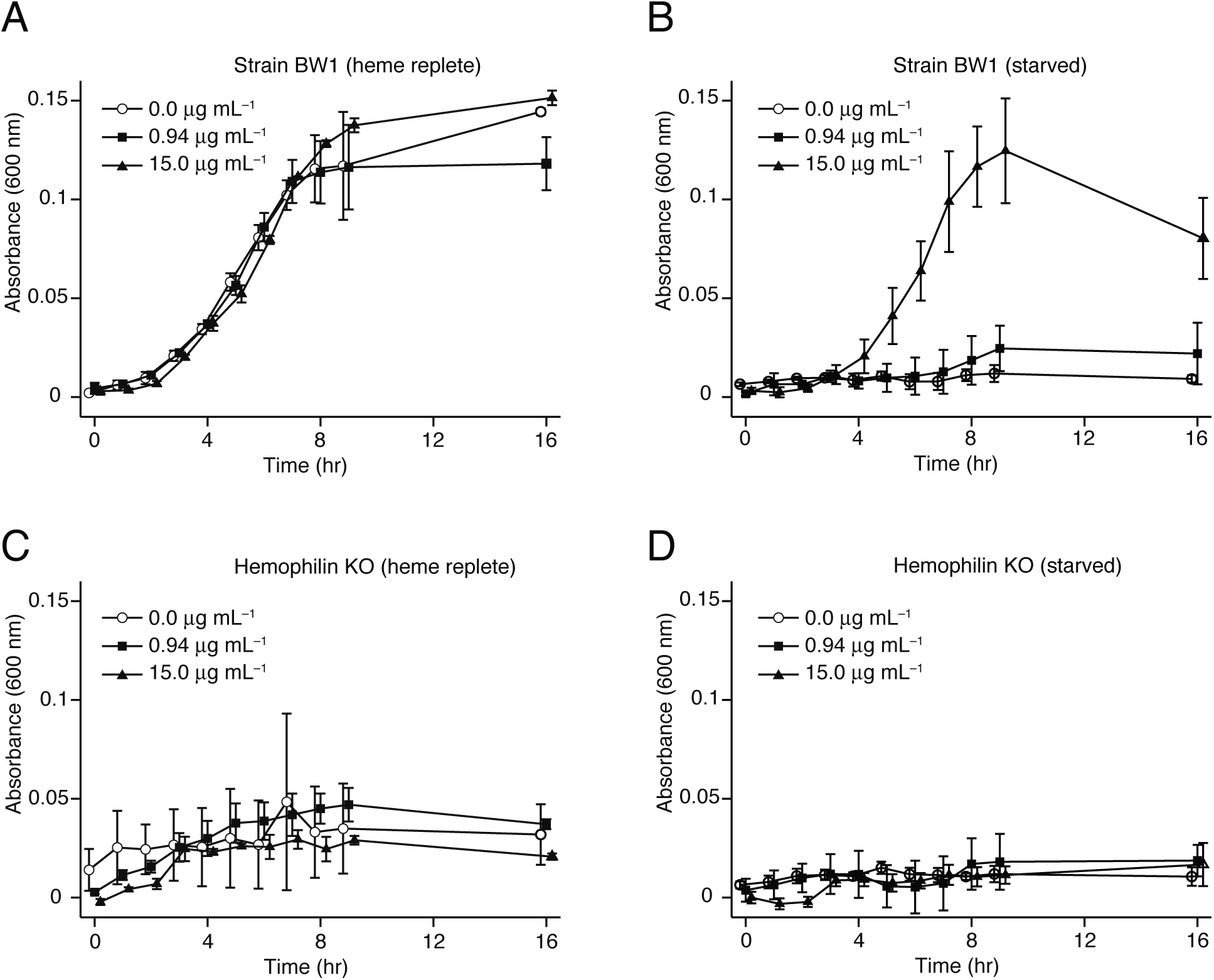
A hemophilin gene knockout strain of BW1 has defects in heme utilisation. Heme-replete and heme-starved populations of *H. haemolyticus* strain BW1 (A, B) or the hemophilin knockout strain (C, D) were inoculated into medium containing varying concentrations of heme as indicated. Error bars represent ±1 SD (n=3).

To test directly if hemophilin can deliver heme to *H. haemolyticus*, we grew cultures supplemented with heme or holo hemophilin at matching final molar concentrations. When BW1 was initially cultured with growth-limiting concentrations of heme (0.6 μg mL^−1^), subsequent addition of holo hemophilin resulted in approximately 3-fold increase in culture density at 20 hours (Supplementary Fig. 15). This growth was the same as growth on an equivalent amount of free heme (Supplementary Fig. 15), suggesting that hemophilin-bound heme was readily available to BW1. In contrast, the addition of holo hemophilin to NTHi caused a decrease in growth (approximately 2-fold decrease in cell density at 20 hours; Supplmentary Fig. 15). Moreover, we found that the inhibition of NTHi in the agar well diffusion assay was overcome by addition of excess heme (Supplementary Fig. 16). Together, these results suggest that hemophilin binds heme in a form that can be utilised by *H. haemolyticus* BW1, but is not accessible to NTHi, and that inhibition of NTHi occurs by heme starvation.

### Distribution of hemophilin across *H. haemolyticus* strains

To gain insight into the broader biological significance of hemophilin, we searched the public sequence databases. From 46 complete or draft *H. haemolyticus* genomes available in Genbank, 30 genomes contained an ORF with 67–100% nucleotide sequence identity to BW1 hemophilin (Supplementary Table 4). In addition, 18 *H. influenzae* genomes (from a total of 700 genome assemblies), 3 *H. quentini* genomes (from a total of 3 assemblies), 1 *H. parainfluenzae* genome (from a total of 40 assemblies) encoded an ORF with similarity to BW1 hemophilin. In each case, the hemophilin gene was immediately downstream of a predicted TonB dependent transporter (Supplementary Table 4). Across the 25 unique hemophilin-like sequences encoded by all genomes, amino acid differences overwhelmingly mapped to surface exposed sites and surface loops on the hemophilin structure (Supplementary Fig. 17). Residues surrounding the heme ligand, including the heme coordinating His119 and the distal pocket residues Gln74, Met79 and Arg82 were 100% conserved in all hemophilin variants, indicating that heme binding is preserved in all variants. In summary, hemophilin homologues are present in a large proportion (63%) of *H. haemolyticus* genomes, compared to a small minority (<3%) of *H. influenzae* genomes, making hemophilin a distinctive feature of *H. haemolyticus* (and also *H. quentini*). Biochemical and genetic studies are now needed to establish if/how hemophilin alleles from the different phylogentic groups contribute to heme uptake in these strains.

The observation that over half of published *H. haemolyticus* genomes carry a hemophilin-like gene raises the question of why only two NTHi-inhibitory strains (BW1 and RHH122) were identified in our functional screen. To start to investigate this question, we searched our collection of 100 *H. haemolyticus* clinical isolates for ORFs that were highly similar to BW1 hemophilin by real-time PCR and DNA sequencing (Supplementary Table 5). By real-time PCR, 15 isolates were positive; this primer set would theoretically have produced an amplicon for 7 of the 46 public *H. haemolyticus* genomes; the same 15% hit rate. We found 6 isolates, including BW1 and RHH122, with ORFs that were 100% identical, and found that the NTHi-inhibitory activity recovered from these isolates varied considerably (Supplementary Table 5). This suggests that high-level production of hemophilin is an unsual feature of BW1/RHH122 and emphasises the importance of understanding how hemophilin expression is regulated, in addition to the effects of sequence variation within the hemophilin protein.

## Discussion

### Hemophilin is a previously undescribed hemophore in H. haemolyticus

In a screen for potential probiotic strains of *H. haemolyticus* that produce inhibitors of NTHi, we identified hemophilin, a small soluble heme binding protein that is secreted at high levels by two *H. haemolyticus* isolates, BW1 and RHH122. Our data suggest that hemophilin plays a positive role in heme acquisition in *H. haemolyticus*, and that the inhibition of NTHi, a pathogenic microorganism that shares a similar ecological niche in the upper respiratory tract of humans, is likely to involve competition for nutrient heme.

*H. influenzae* and *H. haemolyticus* lack the enzymes for *de novo* porphyrin synthesis and depend for their survival on scavenging heme, or protoporphyrin IX plus iron, from the host (Norskov-Lauritsen, 2014). *H. influenzae* have multiple pathways to scavenge heme (Hariadi et al., 2015) (Supplementary Table 4); in each pathway the initial heme binding step is performed by a membrane anchored protein or TonB-dependent outer membrane heme transporter. By comparison, hemophilin represents a previously unidentified mechanism in *Haemophilus* spp—that is, secretion of a diffusible heme binding protein (a hemophore) into the surrounding environment. Hemophores are found in a subset of bacterial species. In Gram-positive species, hemophores can be recognised by the prescence of one or more heme-binding NEAT (near iron transporter) domains; these proteins are covalently attached to the cell surface peptidoglycan, or S-layer, or secreted into the environment (Mazmanian et al., 2000; Grigg et al., 2007; Maresso et al., 2008; Tarlovsky et al., 2010; Malmirchegini et al., 2014). In Gram-negative organisms, three hemophores have been described: HasA from *Serratia marcesens*, *Pseudomonas* spp., and *Yersinia* spp. (Ghigo et al., 1997; Izadi et al., 1997; Ochsner et al., 2000; Rossi et al., 2001); HmuY from *Porphyromonas gingivalis* (Wojtowicz et al., 2009a); and HusA from *P. gingivalis* (Gao et al., 2018). Hemophilin, HasA, HmuY, HusA and NEAT domains have different folds, suggesting that hemophore functions have been independently acquired on multiple occasions in evolution.

Heme coordination by a single His side chain, as seen in hemophilin, is unusual for extracellular heme transport proteins, although it is common for heme proteins in general. Known hemophores make 5-coordinate heme complexes through Tyr (Grigg et al., 2007; Kumar et al., 2013; Kanadani et al., 2015) or 6-coordinate complexes via His/His (Wojtowicz et al., 2009a), Tyr/His (Arnoux et al., 1999), Tyr/Met (Gaudin et al., 2011), or Met/Met (Ran et al., 2007) ligands. Ligand combinations that include Tyr strongly favour binding to ferric over ferrous heme (Reedy et al., 2008), which is appropriate in the typically oxidising environment of the extracellular milieu. Heme ligation through a His ligand would potentially allow hemophilin to capture ferric or ferrous heme, as may occur in aerobic or near anaerobic extracellular environments; similar activity has been attributed to hemophores with His/His (Wojtowicz et al., 2009b) or Met/Met (Nygaard et al., 2006) ligands.

### The heme-site structure suggests hemophilin could bind small molecule ligands in vivo

Notably, hemophilin forms stable complexes with ferrous and ferric heme ligands (Supplementary Figs. 4, 5, 12). The fact that hemophilin purifies from *E. coli* with an O_2_ ligand suggests a potentially biologically meaningful O_2_ affinity and resistance to autooxidation. The exclusion of water from the proximal face of the heme and the small size and enclosed nature of the distal pocket are consistent with this (Carver et al., 1992). Such ligand binding has not been reported for other hemophores, although CO-adducts of the HasA hemophore can be prepared by reduction under pure CO atmosphere (Lukat-Rodgers et al., 2008; Ozaki et al., 2014). The chloride ion present as a heme ligand in the hemophilin crystal structure is not likely to be of physiological importance but indicates a strong propensity to bind to anionic ligands in the ferric state. A chloride ligand has been observed in crystal structures of only four heme proteins, including hemophilin (Kuwada et al., 2011; Singh et al., 2012; Kumar et al., 2013); these proteins have unrelated folds, but all have a similarly positioned Arg side chain in the distal heme pocket (Supplementary Fig. 12). Numerous bacteria heme proteins that act as gas sensors have been described (Martinkova et al., 2013), and because small ligands such as CO, NO and HS^−^ have signalling roles at the host-bacteria interface (Toliver-Kinsky et al., 2019), ligand binding by hemophilin could be biologically important, although this remains to be investigated.

### Proteins with structural similarity to hemophilin bind diverse ligands

The small group of proteins that share structural similarity with hemophilin includes hemoglobin-haptoglobin utilisation protein (HpuA) (Wong et al., 2015), complement factor H binding protein (fHbp) (Schneider et al., 2009) and *Neisseria* heparin binding antigen (NHBA) (Wong et al., 2015), found in members of the *Neisseriaceae*, and transferrin binding protein B (TbpB) (Moraes et al., 2009), which is found in *Neisseriaceae* and some members of the *Pasteurellaceae*, including *H. influenzae* and *H. haemolyticus* (identifiable by blast search). HpuA, fHbp, HNBA and TbpB are lipid anchored in the bacterial outer membrane and have functions in iron uptake, immune evasion or surface attachment. Remarkably, the precise combination of β-barrel topology (8 strands in a meander topology with a shear value of 8) together with hydrophobic residues packed in the barrel core seems to occur only in this group of bacterial proteins (Supplementary Fig. 6).

The hemophores HasA, HmuY and HusA, as well as HpuA and TbpB, deliver cargo to their respective TonB-dependent outer membrane heme transporters (HasR, HmuR, HusR, HpuB and TbpA) (Krieg et al., 2009; Noinaj et al., 2012) that are expressed from the same operon. It was therefore expected that a gene encoding a transporter should be found at a locus close to hemophilin and, indeed, a predicted TonB-dependent transporter with previously uncharacterised function occurs immediately upstream of the hemophilin gene in all 52 published genomes containing a hemophilin gene sequence (Supplementary Table 4). Molecular modelling using the phyre and I-TASSER web servers confidently predicts that the putative hemophilin receptor has a 22-strand β-barrel and plug structure with similarity to TbpA and HasR (Supplementary Fig. 18).

### *Hemophilin* genes are prevalent amongst strains of *H. haemolyticus*, but not *H. influenzae*

A question arises as to why hemophilin-like genes are common in *H. haemolyticus*, yet extremely rare in *H. influenzae*, given the similar requirement for heme and the close phylogenetic relationship between these species. Part of the answer might lie in the different combinations of heme uptake genes in *H. haemolyticus* and *H. influenzae* genomes. *H. influenzae*, and particularly NTHi, have enormous genetic diversity due to their intrinsic transformability and the high rate of recombination with exogenous DNA from their environment (Mell et al., 2011), such that less than 50% of ORFs are present in all strains (the core genome) while the remainder are present variably and represent an accessory genome (Garmendia et al., 2012; De Chiara et al., 2014). In a group of 88 *H. influenzae* genome assemblies (NTHi and Hib) (Pinto et al., 2018), 100% of strains have genes for the outer membrane heme transporters, *hup* and *hemR*, as well as the *hxuCBA* genes for heme uptake from hemopexin (Supplementary Table 4). In addition, 98% of strains have at least 1 gene encoding a transporter for heme uptake from hemoglobin/haptoglobin (*hgpA, hgpB, hgpC*). Thus, the majority of these *H. influenzae* strains have highly redundant pathways for accessing heme from a variety of host sources. The number of heme acquisition genes found in *H. haemolyticus* strains is much more variable. From the 46 available *H. haemolyticus* assemblies, 70% have *hup*, only 7% have *hemR*, and 87% have at least 1 gene for *hgpA*/*hgpB*/*hgpC*. Notably, no *H. haemolyticus* strains carry the *huxCBA* system, which is considered a virulence factor in *H. influenzae* (Morton et al., 2007). Similarly, diagnostic PCR screens performed on large collections of isolates have also shown much lower prevalence of *hup*, *hemR*, and *hxuCBA* genes in *H. haemolyticus* compared to NTHi (Hariadi et al., 2015). On balance then, *H. haemolyticus* have fewer heme uptake pathways than *H. influenzae*, which may have created stronger selective pressure for *H. haemolyticus* to acquire a hemophore. In this context, finding hemophilin-like genes in 63% of *H. haemolyticus* genome assemblies suggests hemophilin may be of considerable importance in the overall heme economy of this species.

### By scavenging heme, hemophilin might be a mechanism of exploitative competition between Haemophilus species

We undertook our initial search for strains of *H. haemolyticus* that could inhibit NTHi with the goal of developing respiratory tract probiotics. The paradigm on which we based this search was that some bacterial strains produce bacteriocins to kill other species that share the same ecological niche (Ghequire et al., 2017; White et al., 2017). The fact that we identified hemophilin, a putative hemophore, suggests that sequestering heme might be another mechanism for bacteria to inhibit their neighbors. Competition for iron (Parrow et al., 2013) and heme (Mozzi et al., 2018) between bacterial pathogens and their hosts is well accepted, and the ability of the host to impose low concentrations of free iron is one of the most important forms of nutritional immunity (Parrow et al., 2013). The emerging picture is that non-pathogenic probiotic bacteria, as well as pathogenic species, have enhanced iron uptake capabilities that facilitate inhibition of microbial pathogens as well as colonisation of the host (Deriu et al., 2013). Our work suggests that competition for heme between heme auxotrophs, such as *Haemophilus* spp, might be similarly important.

Hemophilin is a previously unrecognised heme uptake mechanism in *H. haemolyticus* with the potential to block uptake of essential heme by pathogenic NTHi. Since *H. haemolyticus* co-colonises the upper respiratory tract with NTHi and competes for binding sites on epithelial cells (Pickering et al., 2016), *H. haemolyticus* strains with high-level expression of particular hemophilin alleles, as seen in the BW1 and RHH122, might starve NTHi of heme and locally and specifically inhibit NTHi colonisation.

## Materials and Methods

### Bacterial collection and culture

The origin and method of identification of the bacterial strains has been described previously (Latham et al., 2017). For revival, subculturing, and enumeration of *Haemophilus* spp., chocolate agar (CA) was used and incubated for 18–24 h at 35°C in an atmosphere of 5–10% CO_2_. Isolates were stored at –80°C in 10% w/v sterile skim milk media (SMM).

### Preparation of heme solutions

Solutions (1–5 mg mL^−1^) of ferriprotoporphyrin IX were prepared fresh by dissolution in 0.1 M sodium hydroxide of either bovine hemin chloride (ferriprotporphyrin IX chloride, Frontier Scientific), or porcine hematin (ferriprotoporphyrin IX hydroxide, Sigma-Aldrich) solid, as specified in sections of Materials and Methods. Unless otherwise specified, the term heme is used generically throughout the main text for simplicity, irrespective of the source or oxidation state.

### Agar well diffusion assay

The agar well diffusion assay was performed as described previously (Latham et al., 2017), with the following clarifications. Solid media consisted of 18.5 g L^−1^ brain heart infusion (Oxoid) solidified with 7.5 g L^−1^ Bacteriological Agar (Oxoid), autoclaved at 121°C for 30 min, cooled to 50°C, then supplemented with 1% (v/v) resuspended Vitox® (Oxoid), along with 7.5 mg L^−1^ each of NAD and hematin (Oxoid); these media components are at half the normal concentration typically used for the culture of *Haemophilus influenzae*. 10 mL of agar was dispensed to a 90-mm Petri dish. Indicator strains (NTHi strain NCTC 4560 or NCTC 11315) were prepared by growing for 6–12 h on CA then suspended in Dulbecco’s phosphate buffered saline (DPBS, Gibco) to an absorbance of 1.0, diluting 10-fold with SMM and stored as 100-µL aliquots at –80°C. Thawed aliquots were diluted 100-fold in DPBS and mixed with 5 mL of molten overlay media at a dose predetermined to produce a dense lawn of colonies and immediately poured onto a petri dish of base media. 5-mm diameter circular holes were cut in the agar using a sterile stainless steel cork borer to accept 20–25 µL of test solutions. Plates were left open in a biological safety cabinet for one hour until wells were free of liquid then incubated for 18–24 h at 35°C in a humidified atmosphere containing 5% CO_2_ and the annular radius of cleared zones was recorded. Clearing zone size was affected by media age (older media giving larger the zone sizes), such that treatments were only compared within the same plate.

### Production of native hemophilin

Media for hemophilin production was cold filterable tryptone soya broth (TSB; Oxoid) made to the manufacturer specifications and sterilised by filtration through membrane with a pore size of 0.2 μm, then supplemented with HTM supplement (Oxoid) and Vitox® (Oxoid) according to manufacturers specification. Cultures of BW1 or RHH122 were grown on CA for 12–16 hours, suspended in pre-warmed (37°C) broth in a baffled shakeflask to an absorbance of 0.05 (OD600), then incubated with 200 RPM agitation at 37°C for 24 hours. Culture broths were clarified by centrifugation at 7000 × g for 30 minutes. Hemophilin activity was enriched by ammonium sulfate precipitation at 4°C, with the 50–70% saturation cut collected and redissolved in a volume of PBS equal to 1/20th of the initial culture broth volume then dialysed using a 3500-Da molecular weight cut off (MWCO) dialysis membrane (Thermofisher) for 24 hours at 4°C against 50 mM Tris-HCl, pH 7.5. Following concentration by ultrafiltration with a 10-kDa MWCO centrifugal filter unit (Merck-Millipore) the samples were separated over a Superose 12 HR 10/300 GL column (GE Healthcare) with a bed volume of 24 mL, in 0.15 M sodium phosphate buffer, pH 7.0. Fractions with peak acitvity were applied to a C_4_ reversed-phase HPLC column (Symmetry300; Waters Corporation) that was developed with a linear gradient of 5–95% CH_3_CN, 95–5% H_2_O, 0.1% trifluoroacetic acid. Fractions were collected manually and lyophilised, then resuspended in aqueous buffer of choice.

### RP-HPLC and mass spectrometry

RP-HPLC fractions from the *H. haemolyticus* isolates BW1 or RHH122 that had peak hemophilin activity, or time/volume matched samples from control isolates, were subject to trypsin digestion in batch, or were processed further by Tris-tricine SDS-PAGE, silver staining and in-gel trypsin digestion of individual stained bands, as specified in Results. Silver stain was removed using 30 mM potassium ferricyanide and 100 mM sodium thiosulfate (1:1 mix) and gel pieces were dried by vacuum centrifugation. In-gel trypsin digestion was performed using proteomic grade trypsin (Sigma) as previously described (Wilson et al., 2008). For trypsin digest of HPLC fractions, samples were precipitated in 9 volumes of ethanol, then reduced, alkylated and digested in 100 mM ammonium bicarbonate buffer as previously described (Wilson et al., 2008). Peptide samples were reconstituted in 20 µL HPLC Buffer (2% CH_3_CN, 0.05% trifluoroacetic acid) and analyzed by nanoLC-MS/MS using an Ultimate 3000 RSLCnano HPLC and LTQ-Orbitrap XL fitted with nanospray Flex ion source (ThermoFisher Scientific). Tryptic peptides were loaded at 0.05 mL min^−1^ onto a C_18_ 20 mm × 75 mm PepMap 100 trapping column then separated on an analytical C_18_ 150 mm × 75 mm PepMap 100 column nano-column Peptides were eluted in a gradient from 98% buffer A (0.01% formic acid in water) to 40% buffer B (0.08% formic acid in 80% CH_3_CN and 20% water) followed by washing in 99% buffer B (2 mins) and reequilibration in 98% buffer A for 15 mins. The LTQ-Orbitrap XL was controlled using Xcalibur 2.1 software (ThermoFisher Scientific) and operated in data-dependent acquisition mode where survey scans were acquired in the Orbitrap using a resolving power of 60,000 (at 400 m/z). MS/MS spectra were concurrently acquired in the LTQ mass analyzer on the eight most intense ions from the FT survey scan. Charge state filtering, where unassigned and singly-charged precursor ions were not selected for fragmentation, and dynamic exclusion (repeat count 1, repeat duration 30 sec, exclusion list size 500) were used. Fragmentation conditions in the LTQ were: 35% normalized collision energy, activation q of 0.25, 30-ms activation time and minimum ion selection intensity of 500 counts. The mass spectrometry data have been deposited to the ProteomeXchange Consortium via the PRIDE (Perez-Riverol et al., 2019) partner repository with the dataset identifier PXD013687. Raw MS/MS spectra were searched against a *H. haemolyticus* database downloaded from NCBI on Sept 19^th^ 2016 (37, 881 entries) using the Andromeda search engine in the MaxQuant software (version 1.5.1.2). The settings for protein identification by Orbitrap MS/MS included carbamidomethyl modification of cysteine and variable methionine oxidation, two missed trypsin cleavage allowed, mass error tolerances of 20 ppm then 4.5 ppm for initial and main peptide searches, respectively, 0.5 Da tolerance for fragment ions. A false discovery rate of 0.01 was used for both peptide-spectrum matches and protein identification. PMF analysis of whole RP-HPLC fractions identified EGT80255 peptides as the most abundant ions in samples with NTHi-inhibitory activity obtained from BW1 and RHH122, whereas MS analysis of matched RP-HPLC fractions from control strains, BW29 and BWOCT3, did not identify EGT80255 peptides above background (normalised intensity < 1%). No peptides corresponding to the first 22 amino acids of hemophilin/EGT80255 were identified in gel slices or RP-HPLC fractions. A non-tryptic cleavage site between residues 22 and 23 coincided with a signal peptide cleavage site predicted by signalp 4.1 (Bendtsen et al., 2004). Information from the MaxQuant output is compiled in Supplementary Table 5.

### Recombinant hemophilin

Genomic DNA was extracted from *H. haemolyticus* strain BW1. Primers NIS-F1 (5’-ATTACATATGCAGGTAGTGGGAAATGTATCA-3’) and NIS-R1 (5’-TTATCTCGAGTTAATTTTTAGTACCGCCAAA-3’) were used to amplify a DNA fragment corresponding to the hemophilin ORF residues 23–272 (missing the predicted signal peptide). The hemophilin(23–272) fragment was cloned into the NdeI/XhoI sites of pET28a to express hemophilin with an N-terminal hexahistidine tag. A N-terminal truncated version of hemophilin, hemophilin(55–272), was similarly constructed by PCR cloning with primers NIS-F2 5’-ATTACATATGGACAGTAGTATTCCTAATGAT-3’ and NIS-R1. H89Q and H119Q mutants were generated by standard overlap PCR using the NIS-F1 and NIS-R1 primers together with NIS-89Q-F 5’-TGGATTTCACAGCTTACAGG-3’; NIS-89Q-R 5’-CCTGTAAGCTGTGAAATCC-3’; NIS-119Q-F 5’-GCCAGATCAGCGTGGCTTAGG-3’; NIS-119Q-R 5’-CCTAAGCCACGCTGATCTGGC-3’. Sequenced clones were transformed into *E. coli* strain Rossetta-2 (Novagen), grown in lysogeny broth (LB-Miller) containing 34 μg mL^−1^ chloramphenicol and 25 μg mL^−1^ kanamycin; expression was induced with 1 mM isopropyl β-D-1-thiogalactopyranoside for 3 hours with shaking at 37°C. Hemophilin(23–272) and hemophilin(55– 272) expressed and purified in similar yield; however, the 55–272 deletion variant showed no NTHi-inhibitory activity and was not explored further. For hemophilin produced for crystallography, hemin chloride (5 μM final) was added to expression cultures one hour after induction. Bacterial cell pellets were collected by centrifugation, resuspended in lysis buffer (0.3 M NaCl, 0.05 M sodium phosphate, 0.02 M imidazole, 100 μM phenylmethylsulfonyl fluoride, pH 7.2), lysed by sonication, and clarified by centrifugation. Ni-affinity resin (Gold Biotechnology or Invitrogen) chromatography was performed by gravity at 4°C. The Ni-affinity column was developed with step-wise isocratic gradients increased from 0.02 to 0.25 M imidazole. Peak hemophilin fractions were dialysed against buffer at 4°C (0.3 M NaCl, 0.025 M Tris⋅HCl, 0.02 M imidazole, pH 8.0 at 21°C). The His-tag was cleaved by digestion with thrombin (Sigma-Aldrich) in the presence of 2 mM CaCl_2_ at 37°C for 2 hours; the His-tag was then removed by passing the sample over a second Ni-affinity column. Samples were dialysed at 4°C against 0.02 M sodium phosphate, pH 7, for loading onto cation exchange. Apo (colourless) and holo (orange-red) protein fractions of hemophilin were obtained by cation exchange (UnoS, BioRad), developed with a linear gradient: 0–0.2 M NaCl, 0.02–0.05 M sodium phosphate, pH 7. It was noted that repeated chromatographic separations of the holo protein by Ni-affinity, cation exchange, or SEC (Superose 12 HR 10/300 GL column; GE Healthcare) did not lead to detectible loss of heme, leading us to conclude that apo and holo pools of hemophilin protein were present in *E. coli* during expression and following cell lysis. Heme could be removed from holo hemophilin by acid acetone extraction using the method of Ascoli et al. (Ascoli et al., 1981) or by RP-HPLC (Fig. 2B), to yield apo hemophilin. Apo hemophilin produced by unfolding/reforlding had NTHi-inhibtory and heme binding properties indistinguishable from the properties of apo protein derived from *E. coli* lysates. The concentration of apo hemophilin was determined by spectrophotometry using an extinction coefficient, ε_280_ = 25.9 × 10^3^ M^−1^ cm^−1^ at 280 nm wavelength, calculated from amino acid composition. The concentration of the holo protein was determined using an extinction coefficient, ε_280_ = 38.6 ± 3.6 × 10^3^ M^−1^ cm^−1^ at 280 nm determined from area under the curve analysis of RP-HPLC chromatograms of apo and holo hemophilin performed in triplicate. The ~10% error in concentration measurements for the holo protein were below the detection limit in SDS-PAGE comparison of apo and holo samples (e.g., Fig. 2D).

### BW1 hemophilin Knockout

To confirm the role of hemophilin in generating the anti-NTHi activity of strain BW1, an NIS knockout was constructed using insertional inactivation. A partially assembled WGS of *H. haemolyticus* strain 11P18 (Seq ID: LCTK01000015, contig 00016) was used to acquire sequence flanking the hemophilin ORF for PCR primer design. These PCR primers, NIS-KO-F (gctagacgtgctgatgtt) and NIS-KO-R (tgttgttcttgtcgttgttg) were then used to generate a 1691 bp fragment using genomic DNA from strain BW1 as template. This 1691 bp fragment containing the hemophilin ORF (bp 700–1518) and a unique *Bsp*TI site was cloned in to pGEM-T (Promega) according to the manufacturer’s instructions. An 1132 bp kanamycin resistance cassette, generated by PCR with *Bsp*TI tagged primers Kana-Bsp-F (5’-GCGCCTTAAGTAAACCTGAACCAAA-3’) and Kana-Bsp-R (5’-GCGCCTTAAGGTCGTCAGTCATAAA-3’) using pLS88 (Genbank L23118) as template, was then sub-cloned into the *Bsp*TI site using standard methods. The hemophilin ORF containing the kanamycin cassette was then PCR amplified using NIS-KO-F and NIS-KO-R primers and transformed into strain BW1 using the MIV method (Herriott et al., 1970). Transformants were selected on CA supplemented with 50 mg L^−1^ kanamycin and confirmed by sequencing of the hemophilin region. Sequence verified transformants were tested for the NTHi inhibitory phenotype using an ammonium sulfate extract of a broth culture in a well diffusion assay as previously described (Latham et al., 2017).

### UV-visible spectroscopy

UV-visible spectra were recorded on a Jasco V-630 spectrophotometer fitted with a temperature-controlled sample holder (Jasco) and a septum sealed spectrosil quartz cuvette with a path length of 1.0 cm (Starna, Baulkham Hills, Australia). Samples were prepared at final protein concentration 5– 7 µM in 0.1 M sodium phosphate, pH 7.0, unless otherwise stated. Solutions of heme were prepared fresh by dissolving hemin chloride (Frontier Scientific) to a concentration of ~1 mM in 0.1 M NaOH; for chloride/fluoride binding experiments, hematin was used in place of hemin chloride. Solutions were filtered through a PVDF membrane with a pore size of 0.45 μm (Millipore) and concentrations were determined spectrophotometrically using an extinction coefficient ε_280_ = 58.4 × 10^3^ M^−1^ cm^−1^ in 0.1 M NaOH (Dawson et al., 1969). Ferrous heme was prepared by dilution of heme into nitrogen–purged buffer with addition of molar excess of sodium dithionite; the formation of ferrous heme was monitored spectrophotometrically. All liquid transfer steps were performed under positive N_2_ pressure using a gas-tight syringe (SGE Analytical Science). O_2_ and CO adducts of holo hemophilin were formed by bubbleing O_2_/CO gas. Ferric ligands, CN^−^, HS^−^, Cl^−^, F^−^ were supplied as solutions of KCN, Na_2_S, NaCl and NaF, respectively.

### CD spectropolarimetry

For CD, protein samples were exchanged to 25 mM sodium phosphate, 125 mM NaF, pH 7.0 by SEC over a 24-mL Superose 12 column. Samples were prepared at protein concentration 2.8–3.2 µM in a spectrosil quartz cuvette with a path-length of 0.1 cm (Starna). CD spectra were recorded on a Jasco model J-720 spectropolarimeter with a 450-W water-cooled xenon lamp over the spectral range 260–190 nm; the high tension voltage remained < 450 V. Data acquisition and processing was performed in Spectra Manager Version 1.54.03 with the following acquisition parameters: number of accumulations, 5; bandwidth, 1 nm; response rate, 0.5 s; scan speed 50 nm min^−1^; data pitch 0.5 nm; baseline corrections were performed by subtracting the spectra obtained from matched buffer samples. Prior to sample measurements, the CD amplitude was checked, and if necessary calibrated, using a freshly made solution 1S-(+)-10-camphorsulphonic acid (CSA).

### Crystallography and structure analysis

For crystallization, hemophilin was concentrated to 45 mg mL^−1^ in a buffer comprising 30 mM sodium phosphate, 70 mM NaCl, pH 7.3. Hexagonal crystals with distinct orange-red colour were grown by hanging drop method at 18°C in 0.1 M sodium acetate trihydrate pH 4.5, 2 M ammonium sulfate, by addition of 1 μL of protein solution to 1 μL of mother liquor. Crystals were cryoprotected in 25% glycerol before being flash cooled in liquid nitrogen. Diffraction data sets were collected at the Macromolecular Crystallography MX2 beamline, Australian Synchrotron (Clayton, Australia) (McPhillips et al., 2002) at a temperature of 100 K using x-ray wavelengths of 1.45866 Å to a resolution of 2.1 Å and 0.95372 Å to a resolution of 1.6 Å. Crystal diffraction images were processed in xds (Kabsch, 2010), and data were indexed, scaled and merged using the program aimless in the CCP4 package (Winn et al., 2011). Diffraction data collected at 1.45866 Å (8.49986 keV) were phased by single-wavelength anomalous dispersion (Hendrickson and Teeter, 1981) from the heme iron (above the K-edge, which is 7.1120 KeV) using the phaser SAD pipeline in CCP4, with solvent density modification implemented using parrot (McCoy et al., 2007; Cowtan, 2010). Automated model building was performed using the buccaneer pipeline (Cowtan, 2012) in CCP4, followed by iterative rounds of manual building in coot (Emsley and Cowtan, 2004) and refinement with REFMAC5 (Murshudov et al., 2011). The heavy heme iron atom was subject to anisotropic *B*-factor refinement; all other atoms were refinemened with isotropic *B*-factors. Peaks in the anomalous maps were identified at the following atom positions, in order of intensity (peak height/r.m.s): heme Fe (138 σ), Met171 S (23.4 σ), Met236 S (23.4 σ), Fe-coordinating Cl (19.8 σ), Met61 S (18.4 σ), Met98 S (16.7 σ), Met4 S (10.7 σ), Met91 S (10.7 σ), SO_4_ (8.8 σ), SO_4_ (8.7 σ), Met88 S (9.3 σ), SO4 (8.8 σ), Cl (7.6 σ). The anomalous scattering coefficient, f", for Fe, Cl, S at 8.5 KeV is 2.9, 0.63 and 0.50, respectively. Initial models built with the low-resolution data and reflections for *R*_free_ calculation (5% of data) were transferred as input for refinement against the 1.6-Å data in refmac5. Error estimates for bond length measurements around the heme iron were calculated from Cruickshanks diffraction-data precision indicator (Cruickshank, 1999) multiplied by 2^1/2^ to convert from average coordinate error to bond length error. The volume of the heme-binding cavity was calculated by castp (Tian et al., 2018) and solvent accessible areas were determined using naccess (Hubbard and Thornton, 1993), based on a 1.4 Å probe radius.

### Propagation of *H. haemolyticus* in growth-limiting heme conditions

Isolates were propagated from SMM stock, followed by two overnight passages on CA at 35°C with 5-10% CO_2_ prior to experimental manipulation. Exposure to non-growth conditions was minimised by maintaining suspensions and diluents at 37°C in heat block with sand or benchtop incubator. A suspension (~1.0 OD_600_) of BW1 and BW1-KNOCKOUT was made in TSB from 8-10 h growth on chocolate agar. This suspension was diluted 1:10 in 5mL pre-warmed TSB supplemented (sTSB) with 2% (v/v) Vitox® (Oxoid), and either 15 or 0 mg mL^−1^ porcine hematin (Sigma-Aldrich) to generate heme-replete and heme-starved populations, respectively. Broths were incubated for 14 h at 37°C aerobically without shaking. Resultant starved and replete suspensions were centrifuged at 3000g for 10 min at 37°C and resuspended in fresh, pre-warmed TSB to an OD_600_ of 0.5. A 1:10 dilution was made in pre-warmed TSB containing vitox and either 0, 0.94 or 15 mg mL^−1^ porcine hematin and incubated in a benchtop incubater at 37°C and 220 RPM. Colony counts were performed on all suspensions to confirm initial viability. The OD_600_ was measured by aliquoting 100 mL of growth into wells of a 96-well plate (Grenier Bio-One) and measured in a plate reader (Infinate 200 PRO, Tecan Life Sciences).

### Detection and sequencing of hemophilin in clinical isolates

Genbank sequences from *H. haemolyticus* strains M19107 (AFQN01000044.1) and M28486 (CP031238: region 316000-318200) were aligned and used to design RT-PCR primers to detect hemophilin in genomic DNA from our collection of 100 *H. haemolyticus* isolates using a method as previously described (Latham et al., 2015). Primers NIS-F 5’-GGCGTTGAGATATATGACAGTAG-3’ and NIS-R 5’-TGTAAGGTGTGAAATCCATTTATCG-3’ were used to screen for hemophilin and generated a 126 bp amplicon from position 148 to 273 of the ORF. Primers SEQF 5’-AATCCAGTATTAGTTGTTGATGC-3’ and SEQR 5’-CTTGGTTGTTTATTGTTAATGTAG-3’ were used for amplifying and sequencing hemophilin and generated an amplicon of 1056 bp that included regions 237 bp upstream and 223 bp downstream of the ORF.

### Data availability

The mass spectrometry data have been deposited to the ProteomeXchange Consortium via the PRIDE (Perez-Riverol et al., 2019) partner repository with the dataset identifier PXD013687. X-ray crystallography data have been deposited on the Protein Data Bank with pdb accession 6om5.

## Supporting information

Supplementary Tables, Figures and References

## Acknowledgements

This work was funded in part by a grant from the Clifford Craig Foundation, Launceston, Tasmania (Grant #170). This research was undertaken in part using the MX2 beamline at the Australian Synchrotron, part of ANSTO, and made use of the Australian Cancer Research Foundation (ACRF) detector.

