## Supplementary Tables, Figures and References for "A heme-binding protein produced by *Haemophilus haemolyticus* inhibits non-typeable *Haemophilus influenzae*"

**Supplementary Table 1. MaxQuant analysis of HPLC fractions from *H. haemolyticus* isolates BW1, RHH122, BW39 and BWOCT3**

This table is supplied is a separate file: **Supplementary_Table_S1.xlsx**

**Supplementary Table 2. X-ray data collection and refinement statistics.**

| | Hemophilin |
|---|---|
| **Data collection** | |
| X-ray source | Australian Synchrotron MX2 beamline |
| Detector | ADSC Q315r |
| Temperature (K) | 100 |
| Wavelength (Å) | 0.95372 |
| Space group | $P3_221$ |
| *a*, *b*, *c* (Å) | 91.043, 91.043, 97.484 |
| α, β, γ (°) | 90, 90, 120 |
| Resolution range (Å)† | 45.52–1.60 (1.63–1.60) |
| Total observed reflections† | 1241513 (64897) |
| Unique reflections† | 62031 (3086) |
| $R_{merge}$ (%)†,* | 7.1 (79.8) |
| $R_{meas}$ (%)†,* | 7.3 (81.8) |
| $R_{p.i.m}$ (%)†,* | 1.6 (17.8) |
| $<I / \sigma I>$ † | 23.2 (4.0) |
| $CC_{1/2}$ † | 1.000 (0.909) |
| Completeness (%)† | 100.0 (100.0) |
| Multiplicity† | 20.0 (21.0) |
| Overall *B* factor from Wilson plot ($Å^2$)† | 21.9 |
| **Refinement** | |
| Resolution range (Å)‡ | 41.49–1.60 |
| Reflections in working set‡ | 58934 |
| Reflections in test set‡ | 3056 |
| $R_{work}$ (%)‡ | 17.12 |
| $R_{free}$ (%)‡ | 18.45 |
| Cruickshank DPI‡ | 0.059 |
| No. non-hydrogen atoms | |
| Protein | 1985 |
| Ligand/ion | 84 |
| Water | 170 |
| Average *B*-factors per atom ($Å^2$)§ | |
| Protein | 27.24 |
| Heme | 29.85 |
| Heme Cl ligand | 29.98 |
| Other ligand/ion | 38.84 |
| Water | 36.36 |
| R.m.s. deviations‡ | |
| Bond lengths (Å) | 0.008 |
| Bond angles (°) | 1.45 |
| Ramachandran plot¶ | |
| Favoured (%) | 99.6 |
| Allowed (%) | 0.40 |
| MolProbity score¶ | 1.05, $100^{th}$ percentile |
| PDB code | 6om5 |

Values in parentheses are for the outer shell

† Data from AIMLESS

†,* Reported by AIMLESS, relative to the mean of all $I^+$ and $I^-$

‡ Data from REFMAC5

§ Data from BAVERAGE in CCP4

¶ Data from MOLPROBITY

**Supplementary Table 3. A comparison of Fe(III)–Cl bond lengths**

The table shows heme proteins with chloride ligands from the PDB have Fe–Cl distances in the range 2.34–2.69 Å (excluding pdb 4jet in which the Cl atom in the heme pocket is supposed weakly coordinated, or not coordinated). Fe–Cl bond lengths in small molecule crystals are slightly shorter, 2.22–2.30 Å and close to the theoretical ideal.

| Complex | Method [a] | Fe–Cl (Å) | Ref |
|---|---|---|---|
| Hemophilin | xrd | 2.47 | This work |
| *Rj* DypB D153A (pdb 3vec) | xrd | 2.34 | (1) |
| *Rj* DypB D153H (pdb 3ved) | xrd | 2.69 | (1) |
| *Pa* HbV (pdb 3arj) | xrd | 2.40 | (2) |
| *Pa* HbV (pdb 3ark) | xrd | 2.43 | (2) |
| *Pa* HbV (pdb 3arl) | xrd | 2.45 | (2) |
| Yp HasA (pdb 4jet) | xrd | 3.01 [b] | (3) |
| [Fe(PPIX)Cl] [c] | xrd | 2.218 | (4) |
| $[Fe(TpivPP)Cl]^{-}$ [d] | xrd | 2.301 | (5) |
| $[FeCl(OH_2)_5]^{2+}$ | exafs | 2.26 | (6) |
| | | Fe–O (Å) | |
| metMb Fe(III)($OH_2$) pH 9.0 | xrd | 2.11 | (7) |
| metHb Fe(III)($OH_2$) pH 7.1 | xrd | 2.15 | (8) |
| metMb Fe(III)($OH_2$) pH 6.8 | xrd | 2.05 | (9) |

[a] xrd = single crystal x-ray diffraction, exafs = extended x-ray absorption fine structure

[b] The authors propose the Cl is weakly coordinated, or not coordinated, to Fe in this structure

[c] Hemin choride, PPIX = protoporphyrinIX

[d] TpivPP = tetrakis(α,α,α,α-*o*-pivalamido)phenyl porphyrin

**Supplementary Table 4. Distribution of heme uptake genes in Haemophilus spp**

This table is supplied is a separate file: **Supplementary_Table_S4.xlsx**

Only proteins involved in primary capture of heme from the environment in *H. haemolyticus* or *H. influenzae*/NTHi are show in the table; these being either integral outer membrane proteins or secreted proteins. The table contains accession numbers of heme uptake proteins from all 48 *H. haemolyticus* strains for which sequence data was available (as of December 2018; cells highlighted yellow); 18 *H. influenzae* strains encoding a hemophilin-like gene (highlighted green), from a total of 700 *H. influenzae* genomes; and 88 invasive *H. influenzae* strains collected in Portugal over a 24-year period (10) (highlighted blue). Gene/protein sequences were assigned based on BLAST searches.

**Supplementary Table 5. RTPCR screening for hemophilin in a collection of 100 clinical *H. haemolyticus* isolates.**

| Isolate | RTPCR* | Sequencing PCR* | Nucleotide sequence similarity† (%) | Activity‡ | Activity, equalised for culture density‡ |
|---|---|---|---|---|---|
| BW1 | + | + | 816/816 (100) | 5.5 | 3.75 |
| BW5 | + | + | 804/816 (99) | 0 | 0 |
| BW15 | + | + | 816/816 (100) | 2 | 0 |
| BW18 | + | + | 816/816 (100) | 3 | 0 |
| BW36 | + | – | ND | 3 | ND |
| CF14 | + | + | 783/816 (96) | 0 | 0 |
| L19 | + | + | 706/830 (85) | 0 | 0 |
| L117 | + | – | ND | 0 | 0 |
| L152 | + | – | ND | 3 | 0 |
| L153 | + | – | ND | 3 | 0 |
| NF4 | + | + | 816/816 (100) | 2 | 0 |
| NF5 | + | + | 816/816 (100) | 3.5 | 2 |
| NF6 | + | + | 783/816 (96) | 1 | 0 |
| NF11 | + | + | 785/816 (96) | 0 | 0 |
| RHH122 | + | + | 816/816 (100) | 5.5 | 3.5 |

* RTPCR primers NIS-F and NIS-R produced a 126-bp amplicon within the hemophilin ORF; primers SEQF and SEQR produced a 1056 bp amplicon for sequencing; primer sequences are provided in Materials and Methods. Four isolates failed to produce an amplicon with the sequencing primers (presumably due to differences at the primer annealing sites).

† Number of matches/total alignment length

‡ Measured as annular radius of the clearing zone (mm) in the well diffusion assay

ND, not determined

### Supplementary Figures

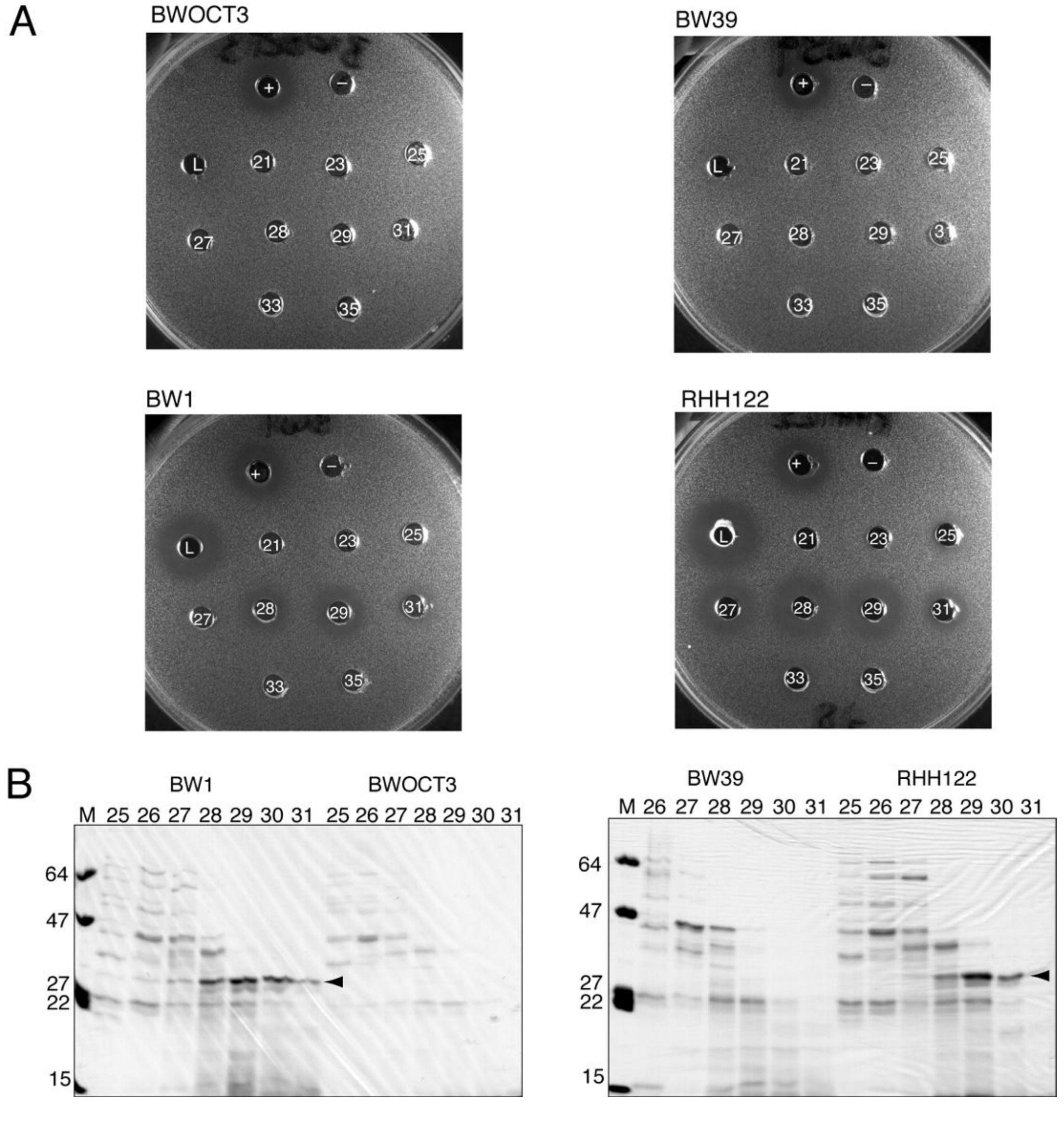


**Supplementary Fig. 1. *H. haemolyticus* isolates BW1 and RHH122 produce an NTHi-inhibitory protein.** Analysis of fractions obtained by SEC separation of ammonium sulfate precipitates of conditioned medium obtained from stationary phase culture of *H. haemolyticus* isolates BW39, RHH122, BW1 and BWOCT3. (A) Agar well diffusion assays of SEC column load (L) and eluted fractions (21–35) for strains as annotated. A control well (+) is concentrated media recovered from a culture of *H. haemolyticus* strain BW1. A negative control (–) is PBS. (B) Tris-glycine buffered SDS-PAGE and Coomassie stain analysis of SEC elution fractions. Fractions with NTHi-inhibitory activity in the agarose well diffusion assay contained a protein of ~30 kDa (arrows) that was absent in matched chromatographic fractions from control strains BW39 and BWOCT3.

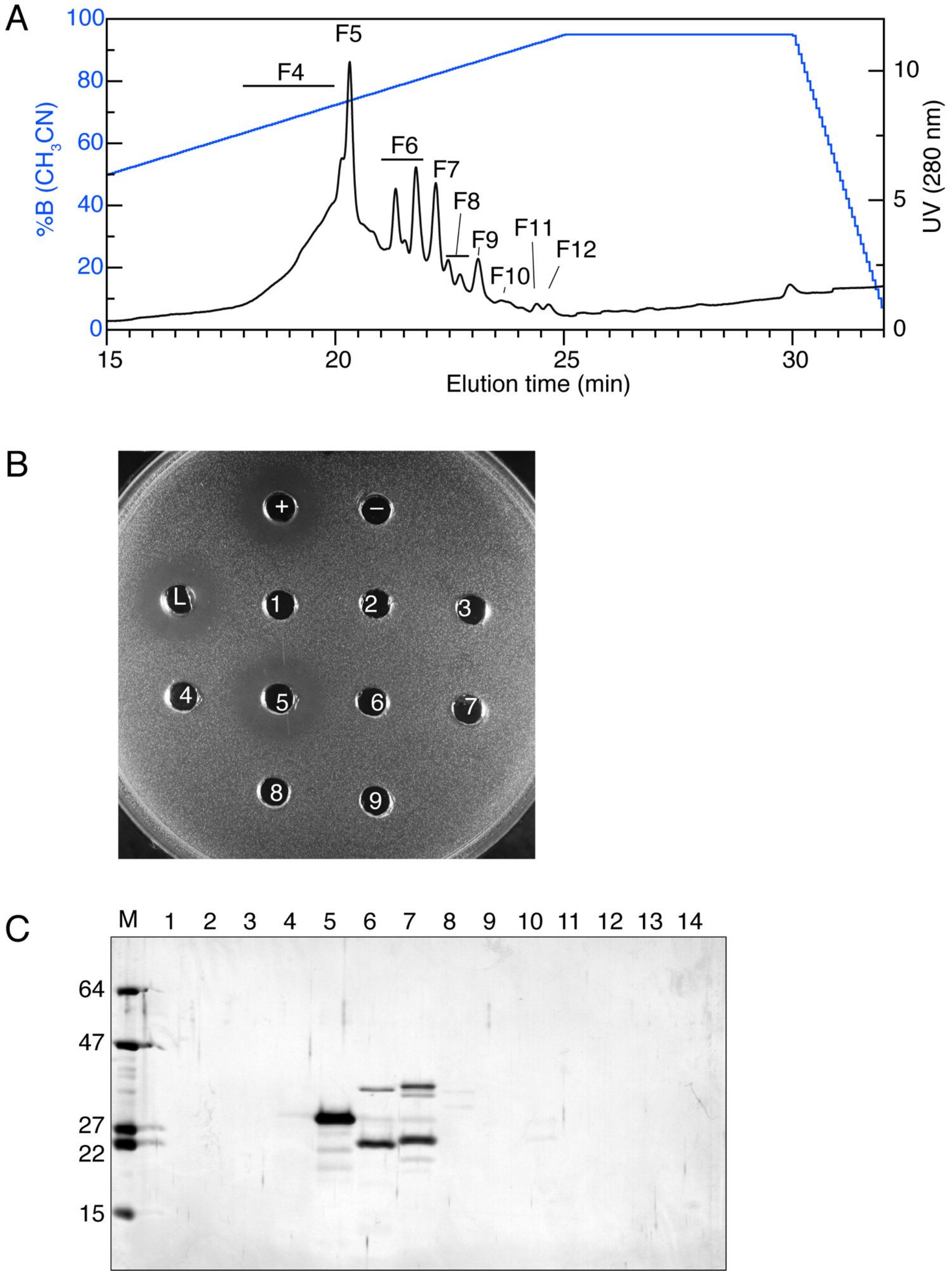


**Supplementary Fig. 2. RP-HPLC purification of hemophilin from *H. haemolyticus* isolate RHH122.** (*A*) Reversed phase HPLC separation of the fraction with peak hemophilin activity obtained from SEC separation of conditioned medium recovered from *H. haemolyticus* strain RH122. (*B*) Agar well diffusion assay of fractions from RP-HPLC separation of RHH122 sample in *A*. NTHi strain 11315 is the indicator strain. Samples from HPLC chromatography load (L) and eluted fractions (1–9) were applied to wells punched in the agar plate. Clear zones around wells indicate inhibition of NTHi 11315. A control (+) is concentrated media recovered from a culture of *H. haemolyticus* strain BW1. A negative control (–) is PBS. (*C*) Tris-tricine SDS-PAGE and silver stain analysis of fractions from HPLC separation of RHH122 sample shown in A, and assayed for activity in B; M, molecular weight markers (kDa).

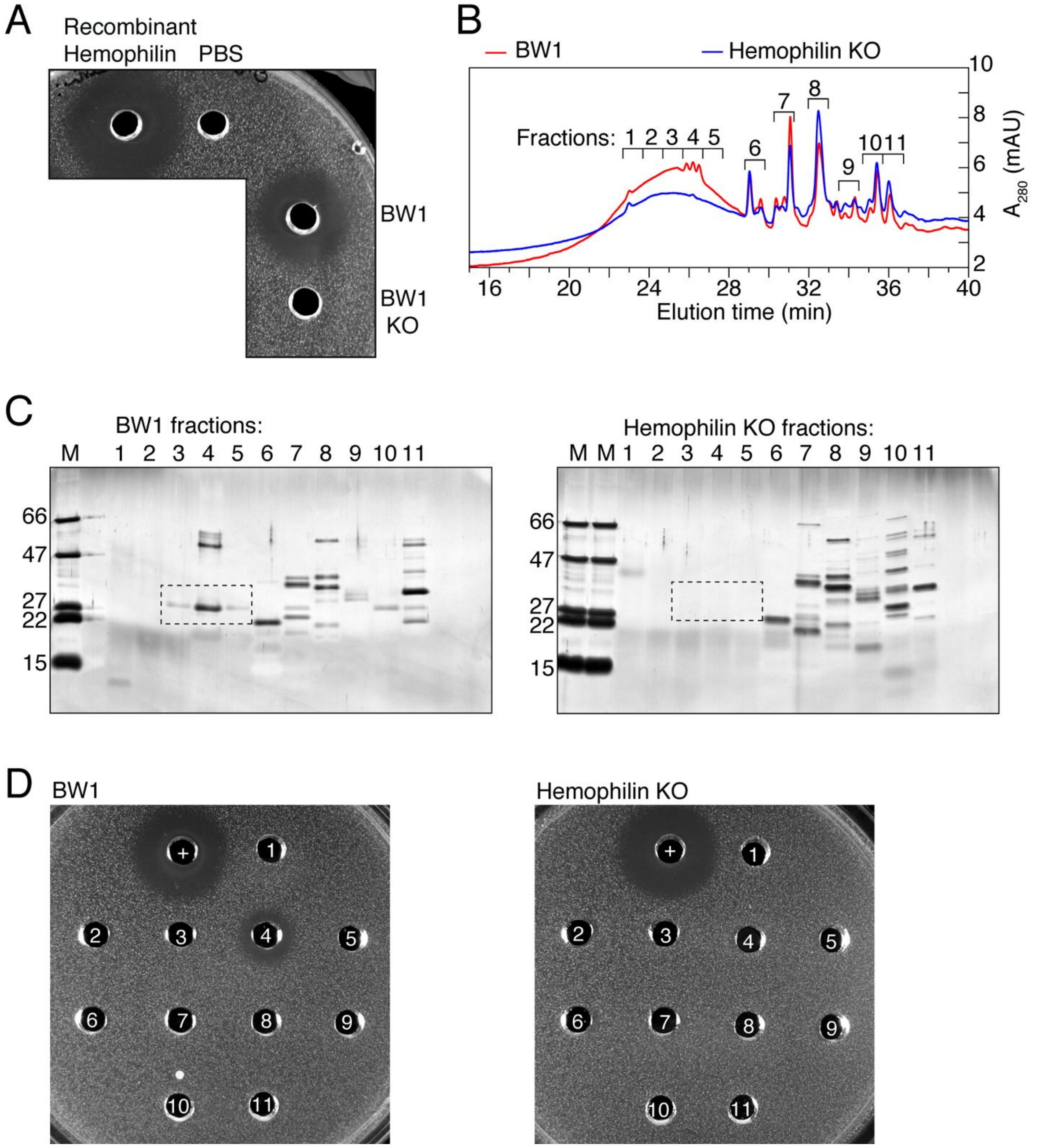


**Supplementary Fig. 3. An *H. haemolyticus* hemophilin gene knockout does not inhibit NHTi.** (*A*) Agarose well diffusion assay: *clockwise from top*, recombinant hemophilin holo protein (400 pmole); phosphate buffered saline (PBS); ammonium sulfate (AS) precipitate from BW1 culture medium equivalent to 1 mL of stationary phase culture at OD 0.88; AS precipitate from the BW1 *hemophilin* deletion strain culture medium equivalent to 1 mL of stationary phase culture at OD 1.1 (BW1 KO). Indicator strain is NTHi strain 11315. (*B*) RP-HPLC of BW1 (red) or hemophilin KO samples (blue) after AS and SEC purification as described in methods. Fractions were collected by hand, as indicated, for further analysis. The size of the hemophilin peak(s) was variable between experiments: compare Fig. S4B with HPLC traces in Figure 1 and fraction 5 of Figure S2 (note, different elution gradients were employed). We attribute this to variable expression level between experiments. Factors that regulate hemophilin expression level are unknown. (*C*) Tris-Tricine SDS-PAGE and silver stain analysis of fractions collected as shown in B. Dashed box indicates the position of the 27-kDa hemophilin band present in BW1 fractions, but not in the hemophilin KO samples; M, molecular weight markers (kDa). (*D*) Agarose well diffusion assays of RP-HPLC samples as annotated in B, C. Activity is detected only in fraction 4 from BW1. The positive control is (+) is purified hemophilin (340 pmole); the negative (–) is PBS buffer.

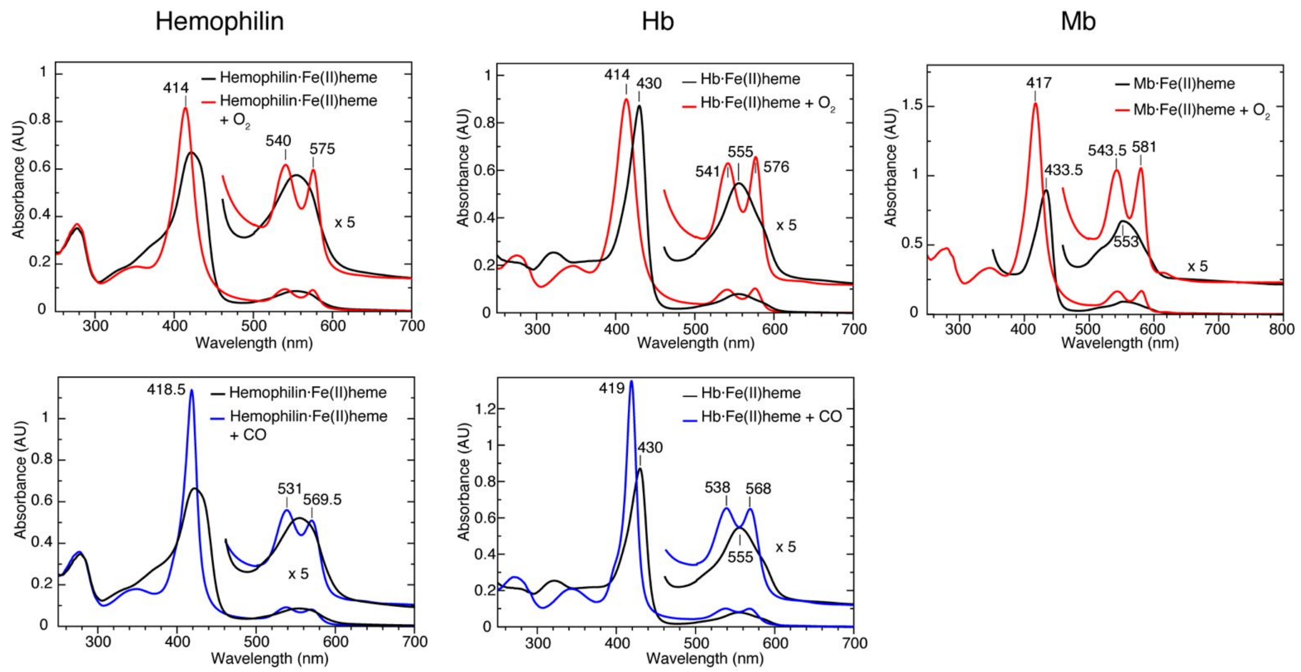


**Supplementary Fig. 4. Hemophilin reconstituted with ferrous heme can bind $O_2$ and CO.** Apo hemophilin (10 μM) was reconstituted with ferrous heme, generated by treating heme with excess sodium dithionite under nitrogen. Addition of $O_2$ (red traces) or CO (blue traces) produced spectra with distinct Soret, α and β absorption bands typical of a low-spin 6-coordinate ferrous heme, as evident from a comparison with the spectra of hemoglobin (Hb) and myoglobin (Mb) $O_2$/CO complexes. All spectra were recorded in 0.2 M sodium phosphate buffer, pH 7.0.

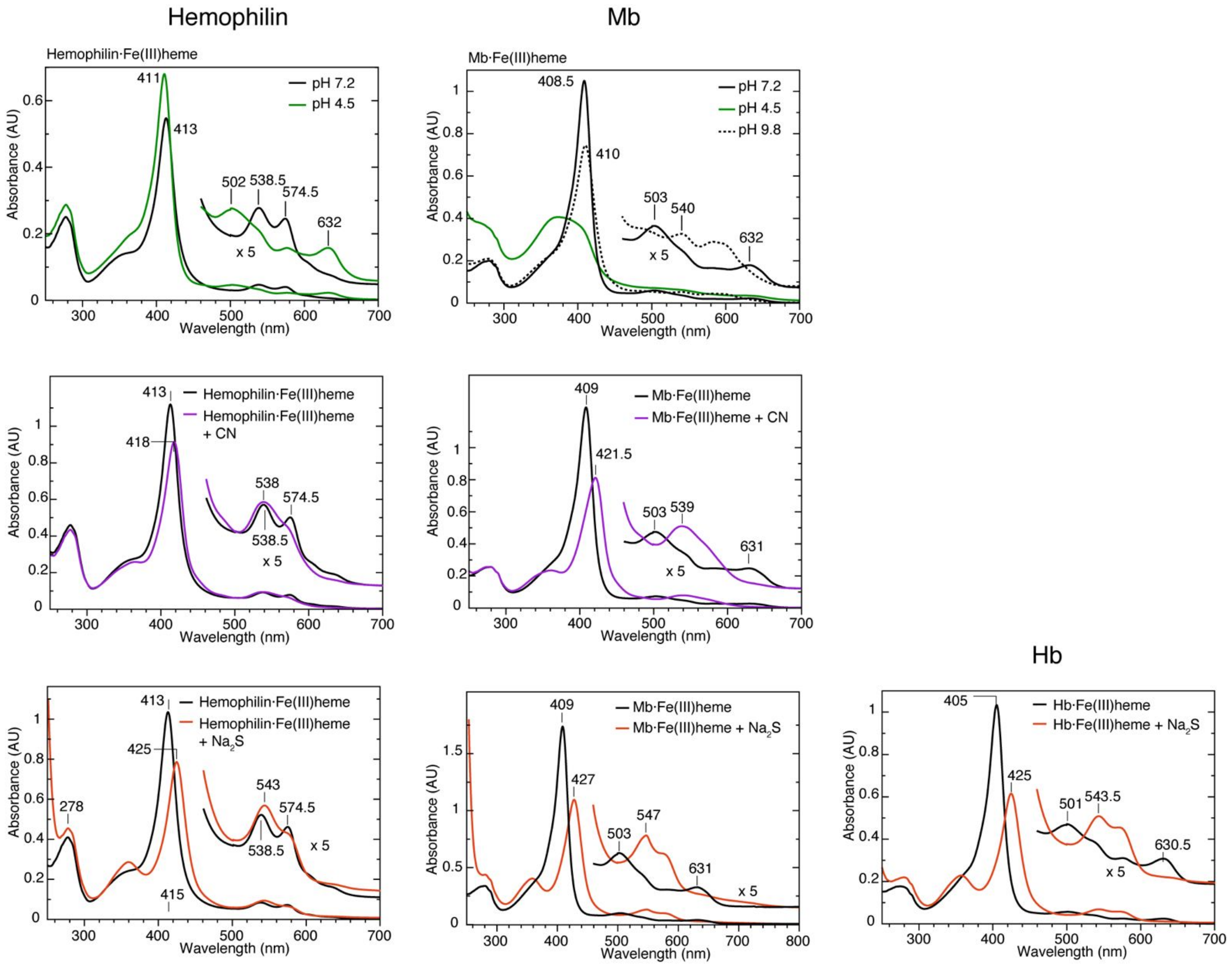


**Supplementary Fig. 5. Hemophilin reconstituted with ferric heme can bind $CN^-$ and $HS^-$.** (*Top*) In 0.2 M sodium phosphate buffer, pH 7.2 (black traces), apo hemophilin (10 μM) reconstituted with ferric heme gave a UV-visible spectrum with α and β absorption bands at 538.5 and 574.5 nm, suggesting a 6-coordinate low-spin Fe(III) with $OH^-$ ligand (20). The aqua complex of Mb at pH 7.2 is high-spin Fe(III) with characteristic charge transfer bands at ~500 nm and 630 nm (20). In 0.1 M sodium acetate, 2 M ammonium sulfate, pH 4.5 (green traces), the condition used for crystallization, hemophilin produces a spectrum that is typical of a high-spin aqua complex. At pH 4.5 Mb is unfolded, but in 0.2 M sodium borate pH 9.8 (dashed trace), Mb converts to a mixture of high spin/low spin hydroxo Fe(III) species. The pKa of the acid alkaline transition of Mb is 8.9 (21, 22). Clearly the pKa of the $H_2O/OH^-$ transition in hemophilin is much lower (pH<7.2). (*Middle, bottom*) Spectra were recorded in 0.2 M sodium phosphate buffer, pH 7.0. Compared to the unliganded form (black traces), addition of KCN (purple traces) or $Na_2S$ (orange traces) to hemophilin produced spectral changes indicating ligand binding, including red-shift in the Soret absorption peak to ~420 nm and the appearance of unresolved α/β bands with absorption maximum ~540 nm, similar to the spectra of $CN^-/HS^-$ complexes of Mb and Hb, which are low-spin 6-coordinate Fe(III) species (20, 23).

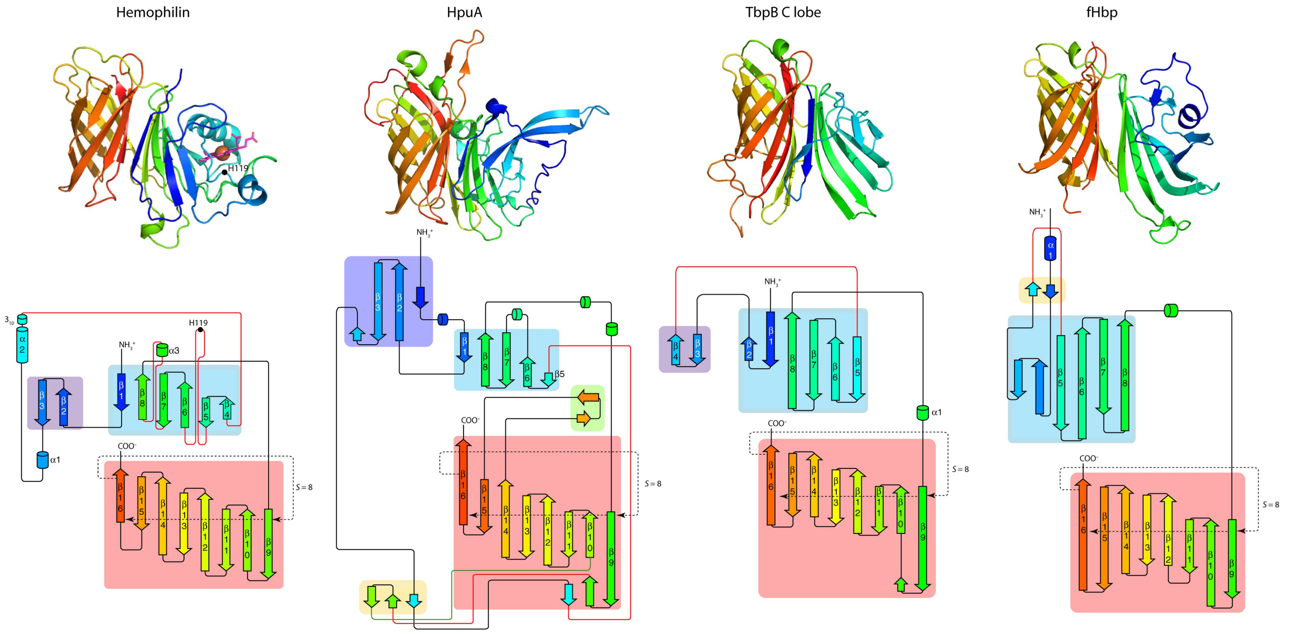


**Supplementary Fig. 6. Structure and topology of hemophilin and similar proteins.** Richardson diagrams (*top*) and topology (*bottom*) for hemophilin and the three proteins with highest similarity to hemophilin according to DALI searches (24), coloured blue through red from N- to C-terminus. These proteins are: hemoglobin-haptoglobin utilisation protein (HpuA) from *Kingella denitrificans* (pdb 5ec6, rmsd 2.4 Å over 185 residues) (25), transferrin binding protein B (TbpB) from *Actinobacillus suis* (pdb 3pqu, rmsd 3.7 Å over 181 residues) (26), and complement factor H binding protein (fHbp; pdb 2w80, rmsd 3.7 Å over 174 residues) from *Neisseria meningitides* (27). A fourth highly similar protein, as identified by DALI, is *Neisseria* heparin binding antigen (NHBA; pdb 2lfu, rmsd 4.0 Å over 120 residues) (28) from *Neisseria meningitides* (not shown). Topology diagrams were produced using PROORIGAMI (29) with manual editing to facilitate comparisons between proteins. Secondary structure elements are numbered to indicate similarity with hemophilin. β-Strands that contribute to the same sheet or barrel structure are enclosed by coloured rectangles. Segments of irregular secondary structure that cross in the diagrams are coloured red/green for unambiguous backbone tracing. The C-terminal β-barrel has an antiparallel meander topology and is an unusual example of an eight-stranded ($n$=8) β-barrel with a shear value of eight ($S$=8) indicating that hydrogen bonding around the β-barrel links residue $i$ and $i$±8 (30). Hemophilin, HpuA, TbpB, fHbp and NHBA (not shown) also share a 2–6 strand β-sheet that packs against the barrel.

Remarkably, the precise combination of β-barrel topology (8 strands in a meander topology with a shear value of 8) together with hydrophobic residues packed in the barrel core seems to occur only in the above group of bacterial proteins. A large number of other proteins contain an 8-stranded β-barrel with meander topology, but with different distributions of polar/non-polar residues (integral membrane *versus* aqueous soluble)(31, 32) and/or shear number (10 or 12)(33-35). Interestingly, the N-terminal region of hemophilin can be interpreted as an opened and distorted 8-stranded β-barrel, with insertions into the loops between the β-3/β-4 strands and between the β-5/β-6 strands forming the docking site for heme, thus providing a hypothetical evolutionary origin for hemophilin through domain duplication and subsequent divergence of an ancestral tandem-barrel structure. We note that the 8-strand β-barrel topology has been adapted to a ligand binding function in other ways, most notably in the lipocalin superfamily, wherein a greater shear ($S$ = 12) produces a barrel with a larger diameter that can accommodate a ligand inside the barrel, for example, a heme group (36) or ferric-siderophore (37).

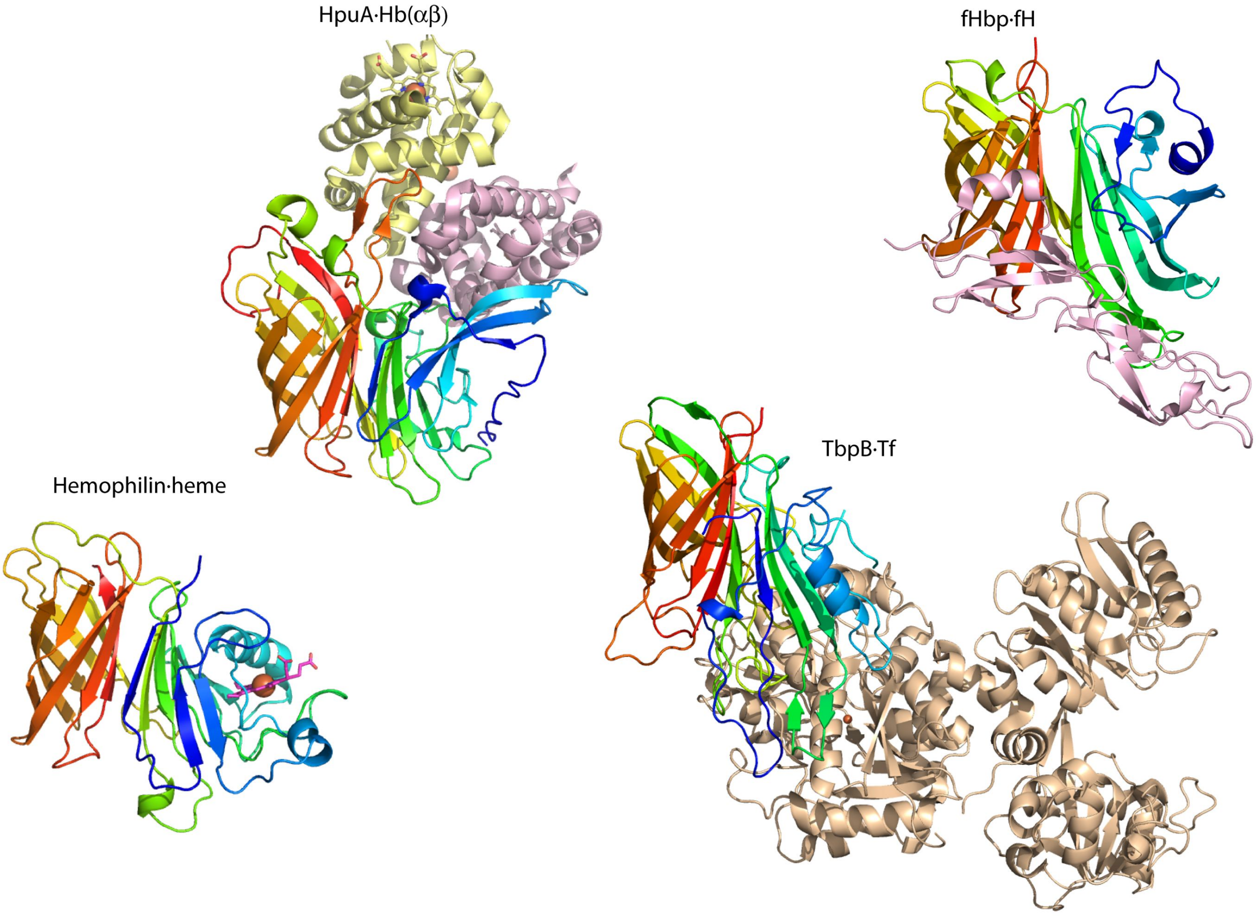


**Supplementary Fig. 7. Different ligands for members of the family of proteins with structural similarity to hemophilin.** Hemophilin is secreted from *H. haemolyticus* strains BW1 and RHH122; HpuA, TbpB, fHbp and NHBA (not shown) are expressed as lipoproteins on the bacterial outer membrane; all have ligand binding functions, but bind very different ligands. The ligand complexes are shown for hemophilin·heme (this work), HpuA in complex with hemoglobin (αβ dimer in the crystallographic asymmetric unit) (HpuA·Hb(αβ); pdb 5ee4) (25), TbpB (N lobe) bound to human transferrin (TbpB·Tf; pdb 3ve1) (38) and fHbp bound to complement control protein (CCP) domains 6–7 of complement factor H (fHbp·fH; pdb 2w80) (27). A different face of the domain engages protein ligand in each case. Note both the N and C lobes of TbpB have structural similarity to hemophilin; the N lobe (shown here) is less similar than the C lobe (shown in Supplementary Figure 6).

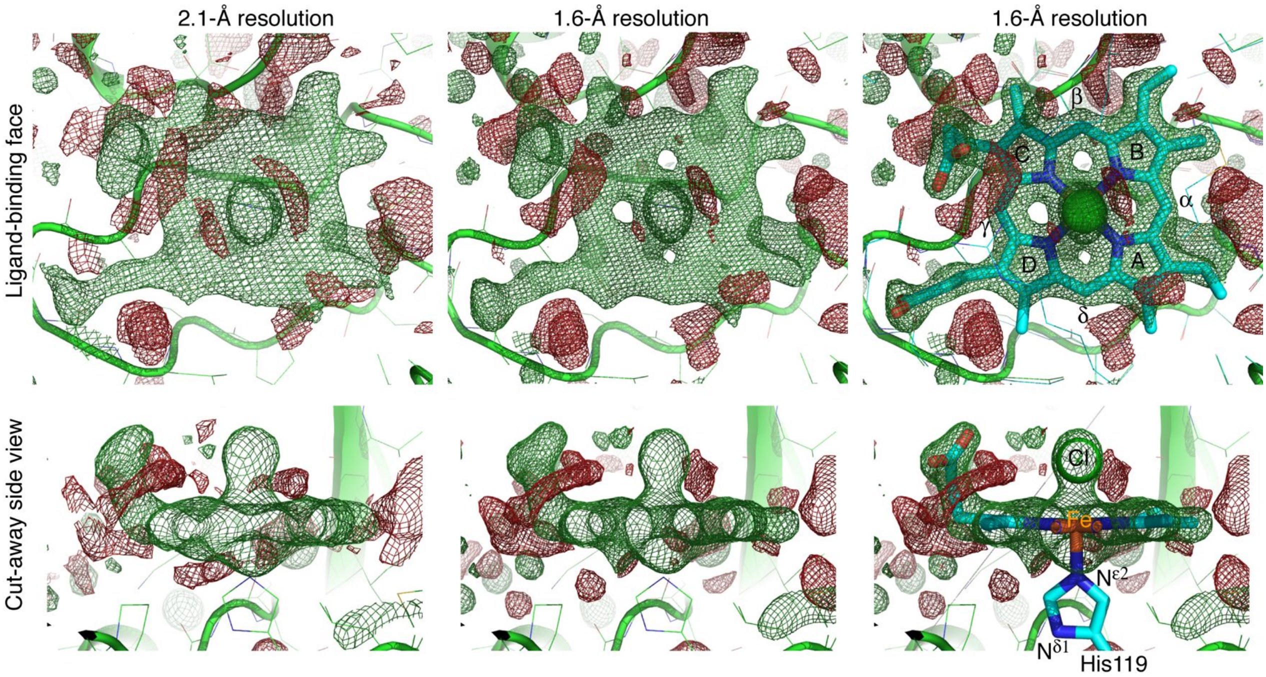


**Supplementary Fig. 8. Identification of heme and a heme-coordinated ligand in electron density, before refinement.** The figure shows portions of $F_O$–$F_C$ electron density maps contoured at +3 σ (green) and –3 σ (red) obtained at intermediate stages of structure refinement, after building all residues of the hemophilin polypeptide, but before before placing ligands, waters or other solvent molecules. (*Left*) Diffraction data collected at an x-ray wavelength of 1.45866 Å to a resolution of 2.1 Å were phased by single-wavelength anomalous dispersion using the PHASER SAD pipeline in CCP4. Automated model building was performed using the BUCCANEER pipeline in CCP4, followed by manual building in COOT to complete the polypeptide (excluding the N-terminal expression tag which was disordered). The Figure shows a portion of the $F_O$–$F_C$ map generated by REFMAC5 that clearly identified the position of a porphyrin with a single-atom ligand, distal to the heme-coordinating His119 side chain. (*Middle*) A data set collected at an x-ray wavelength of 0.95372 Å to a resolution of 1.6 Å was refined against the polypeptide model obtained from the low-resolution data. (*Right*) The figure shows the result of real-space refinement of ferriprotoporphyrin IX and $Cl^-$ ion into the $F_O$–$F_C$ density using COOT. Pyrrole rings and methine bridges of the porphyrin are annotated according to the Fischer system. For reference (not annotated) methyl groups are in the 2,7,12,18-positions, vinyl groups in the 3,8-positions and propionate groups in 13,17-positions, according to the IUAPC-IUB nomenclature of tetrapyrroles for trivial names and locants.

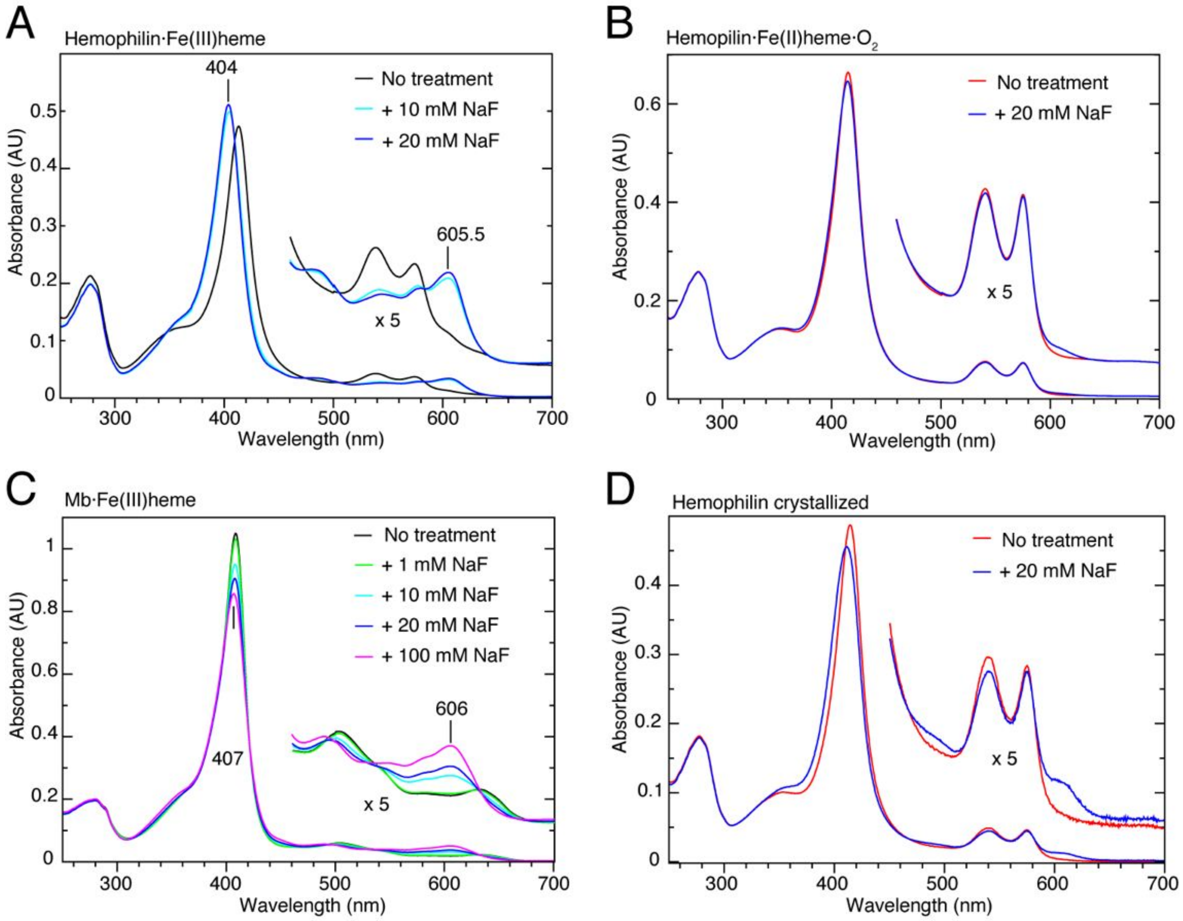


**Supplementary Fig. 9. NaF is a probe for the presence of ferric heme in hemophilin preparations.** As shown in Supplementary Figures 4 and 5, ferrous hemophilin bound to $O_2$ and the ferric hemophilin aqua complex both give UV-visible spectra characteristic of low-spin iron complexes, making it hard to detect low levels of cross-contamination between these forms. To detect contamination of the hemophilin $O_2$ complex with ferric heme we used NaF as follows. (*A*) Large spectral changes, including appearance of a strong charge transfer band at ~606 nm, occur upon addition of NaF to hemophilin·Fe(III)heme (5 μM), consistent with a shift from low spin Fe(III) complex with axial $OH^-$ ligand to a high spin Fe(III) complex with axial $F^-$ ligand. (*B*) A small UV-visible spectral change is observed when NaF (20 mM) is added to hemophilin·Fe(II)heme·$O_2$ (5 μM), freshly purified from *E. coli*, consistent with the presence of ferric heme in a small proportion of hemophilin molecules ($F^-$ being a ligand specific for Fe(III) hemes). (*C*) Binding of $F^-$ to metmyoglobin generates a 6-coordinate high-spin complex similar to that for hemophilin·Fe(III)heme, although higher concentrations of NaF are required suggesting hemophilin has higher affinity for an $F^-$ ligand than does Mb. (*D*) Addition of 20 mM NaF to the hemophilin sample prepared for protein crystallization results in substantial changes in the UV-visible spectrum, including a blue shift of the Soret band and appearance of absorption at ~606 nm, suggesting the presence of contaminating Fe(III) heme. Spectra in A–C were recorded in 0.1 M sodium phosphate buffer, pH 7.0, with addition of NaF as indicated. Spectra in D were recorded in 0.1 M sodium phosphate, pH 7.3 in the absence or presence of 20 mM NaF.

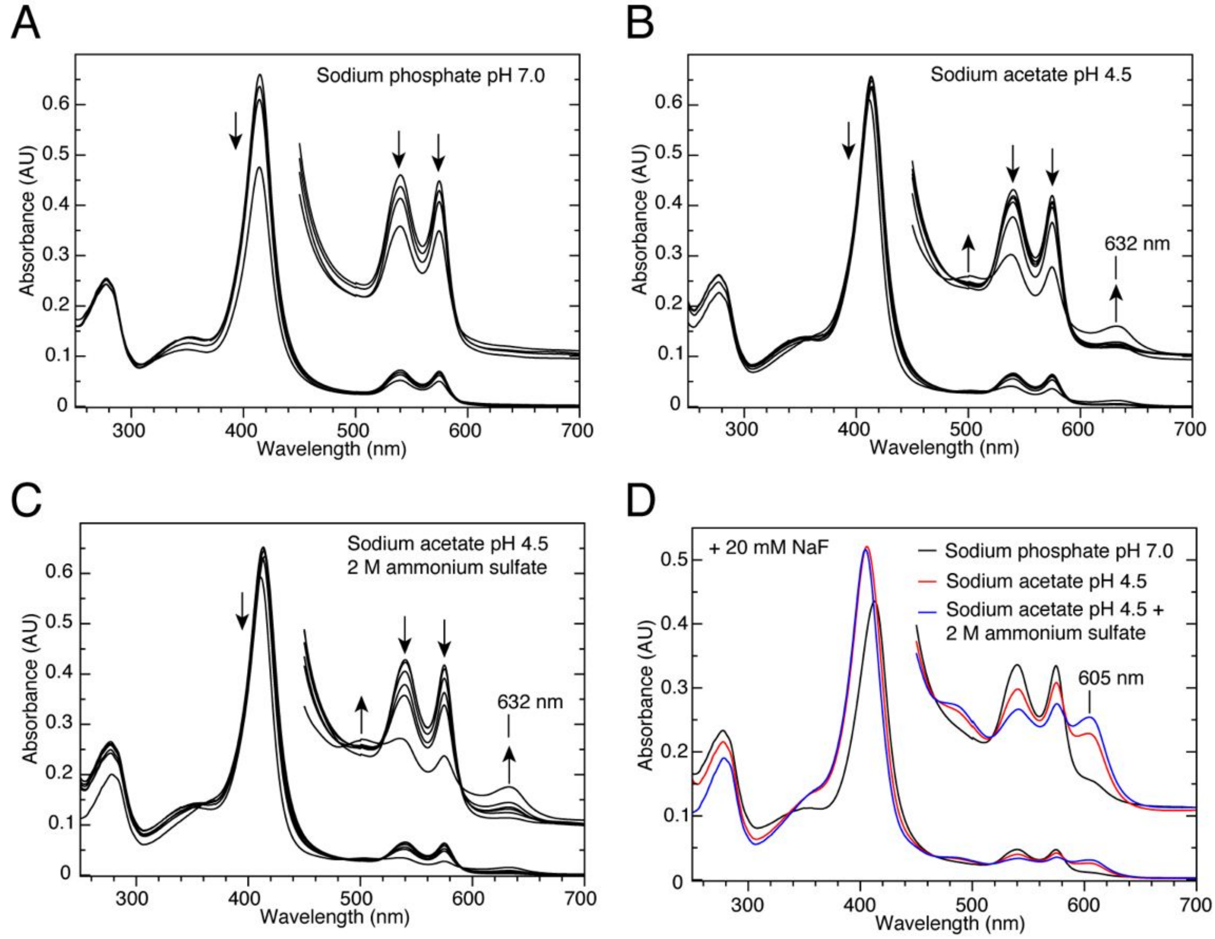


**Supplementary Fig. 10. Stability of hemophilin in crystallization buffers.** (*A*) hemophilin·Fe(II)heme·$O_2$ (6 μM) in 0.1 M sodium phosphate pH 7. Spectra recorded at $t$ = 0, 168, 312, 2256 hours at room temperature. The uniform decrease in spectral intensity across all wavelengths suggests protein precipitation or loss of heme. (*B*) hemophilin·Fe(II)heme·$O_2$ (6 μM) in sodium acetate pH 4.5. Spectra recorded at $t$ = 0, 5, 72, 168, 312, 2256 hours. Appearance of peaks at ~500 and 632 nm indicate the evolution of high-spin Fe(III) heme. (*C*) Conditions as B, with addition of 2 M ammonium sulfate; the condition of the crystallization experiment. (*D*) Samples from the final time point in A–C were adjusted to 20 mM NaF. Formation of a fluoride heme species is indicated by an absorption peak at ~605 nm. A small 605 nm peak at pH 7 confirms that there is little autooxidation at this condition (although there is a decrease in the total soluble hemophilin over time, as shown in A). Both low pH and 2 M ammonium sulfate promote autooxidation.

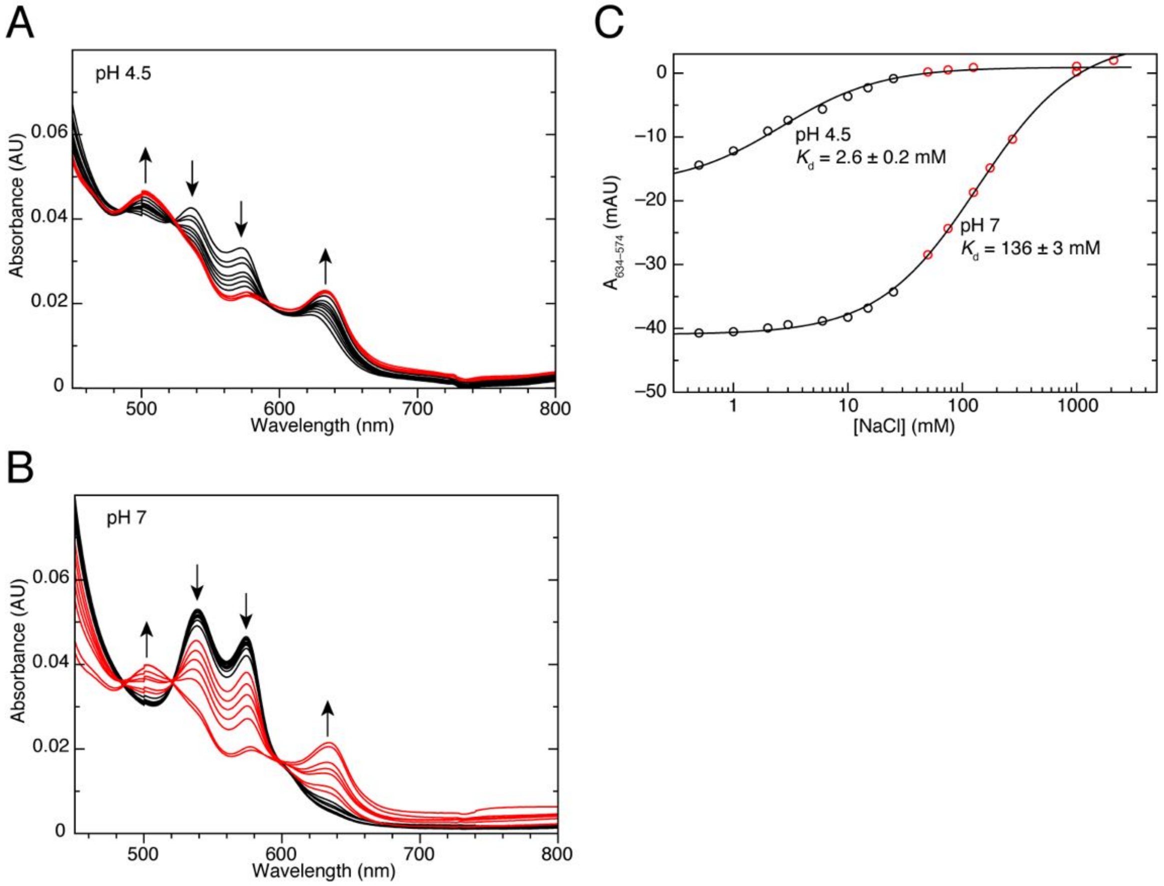


**Supplementary Fig. 11. Ferric hemophilin binds chloride ions.** (*A*) Titration of NaCl into holo hemophilin (6 μM) reconstituted with hematin (ferriprotoporphyrin IX hydroxide) in chloride-free 0.1 M sodium acetate, 2 M ammonium sulfate, pH 4.5. Spectra recorded with [NaCl] in the range 0.5–25 mM are coloured black, and for [NaCl] in the range 50–2000 mM, spectra are coloured red. (*B*) Titration as in A, but with buffer comprising 0.1 M sodium phosphate, pH 7. (*C*) Binding isotherms for chloride calculated from absorbance changes at 574 and 634 nm shown in A, B. Errors are the asymptotic standard errors on the fit parameter $K_d$ (dissociation equilibrium constant) as calculated by GNUPLOT version 4.6.

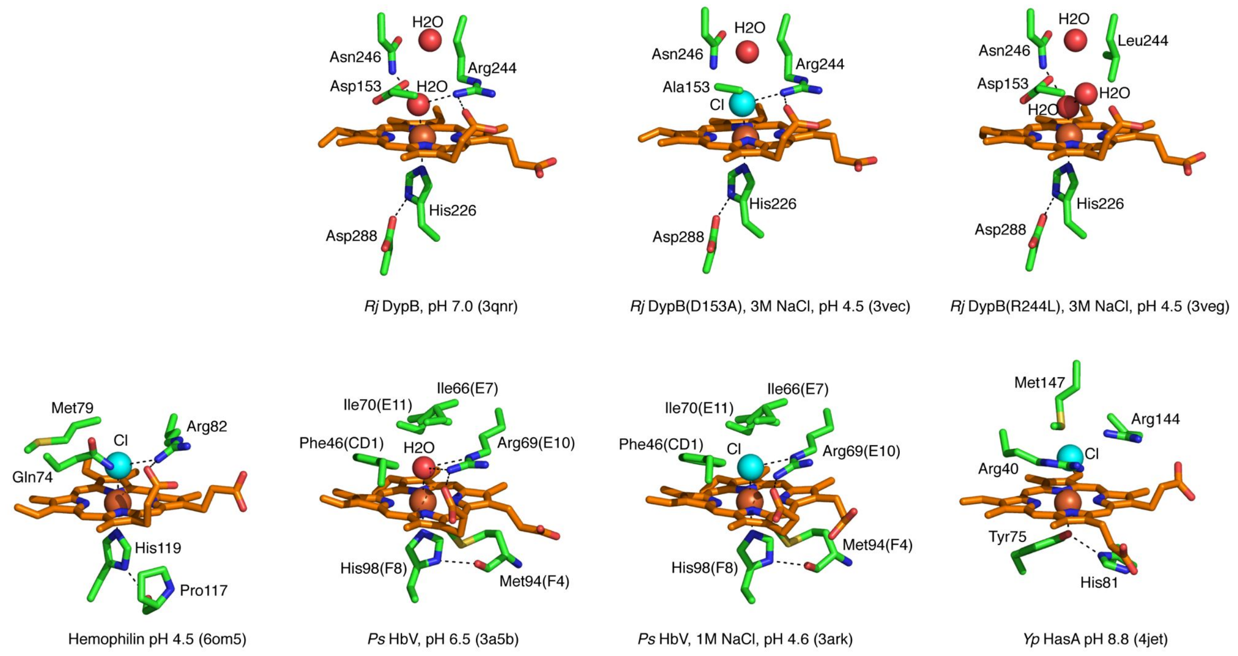


**Supplementary Fig. 12. Heme sites with chloride ligands in the PDB.** Protein crystal structures containing heme with chloride coordinating, or positioned close to, the heme iron have been deposited for three proteins, other than hemophilin. These are the dye decolourizing peroxidase (DypB) from *Rhodococcus jostii* (pdb 3vec, *shown*, and 3ved, *not shown*) (1), the monomeric hemoglobin component V (HbV) from *Propsilocerus akamusi* (pdb 3ark, *shown*, and 3arj and 3arl, *not shown*) (2), and the hemophore, HasA, from *Yersinia pestis* (pdb 4jet) (39). These examples share with hemophilin an Arg sidechain in the distal heme pocket positioned with the guanadinium group roughly planer with the porphyrin ring and within hydrogen bonding distance of the chloride ligand and hydrogen bonding with one of the heme 17-propionate groups. A water ligand can also be accommodated with a similar side chain arrangement in some cases. Mutational and structural analyses have confirmed a positive role for the distal site Arg in binding to chloride (1) and other ionic ligands such as $F^-$, $N_3^-$, $CN^-$, $SCN^-$, $OH^-$ (40-42); similarly it may also facilitate $HS^-$ binding. The heme site structure of hemophilin suggests that binding to ferrous and ferric heme ligands could occur *in vivo*. Bacteria from a variety of environments use heme proteins to sense CO, $O_2$, NO and $HS^-$ (43, 44), and these small ligands also have powerful signalling roles in the mammalian immune system (45-47), making small molecule binding by hemophilin potentially important. In contrast, an enzyme activity for hemophilin seems unlikely based on the heme pocket structure: for example, oxidoreductase heme enzymes such as horseradish peroxidase and cytochrome c peroxidase (48), dye-decolourizing peroxidase (49), and nitric oxide dioxygenase (50) have an acidic Asp or Glu residue hydrogen bonded to the proximal His.

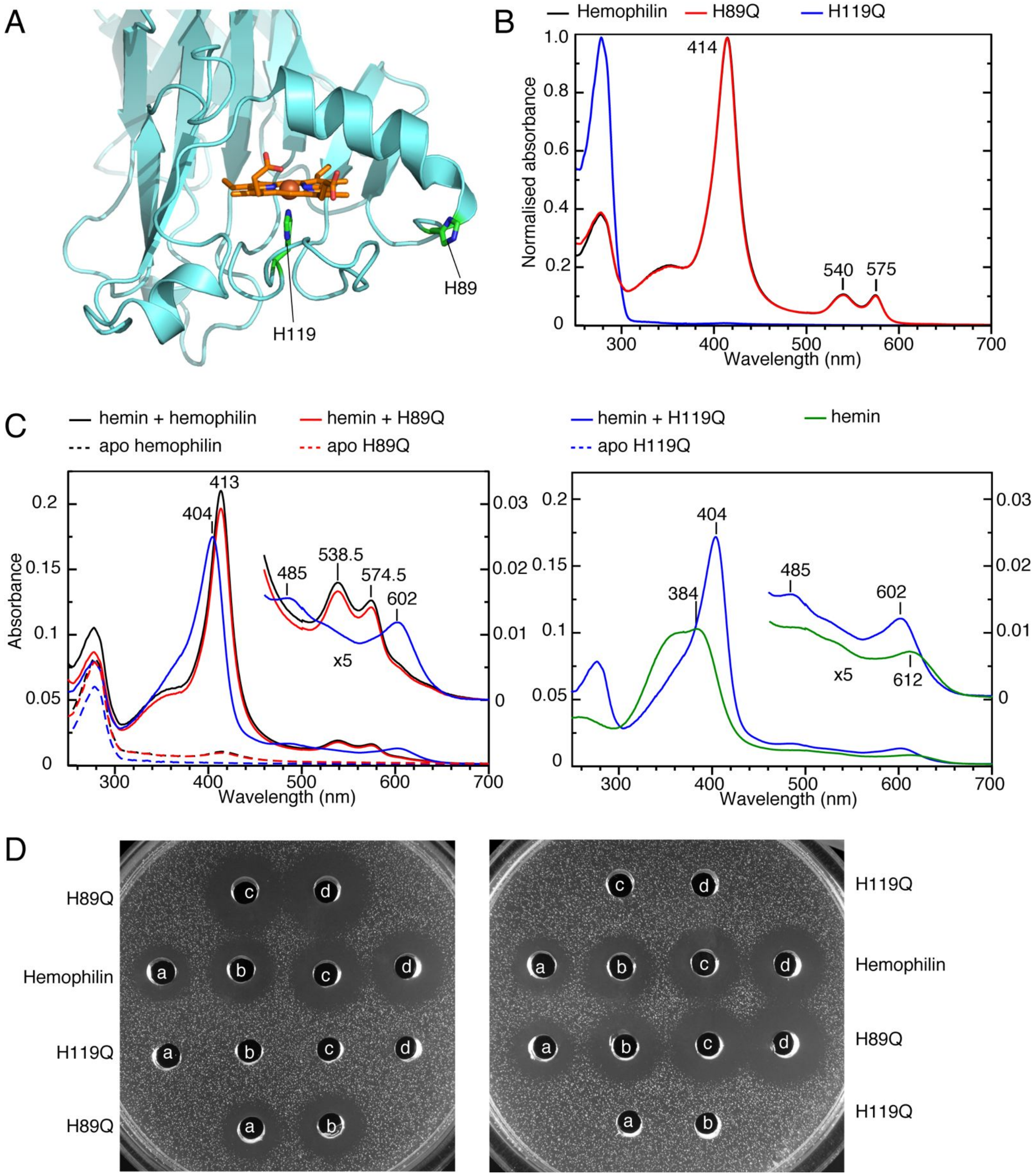


**Supplementary Fig. 13. Activities of hemophilin mutants H89Q and H119Q.** (*A*) Richardson diagram of the N-terminal subdomain of hemophilin showing H89 and H119. (*B*) UV-visible spectra of hemophilin proteins following Ni-affinity purification. Spectra of hemophilin and hemophilin(H89Q) are identical, indicating that the H89Q mutation has no effect on heme binding. The spectrum of hemophilin(H119Q) has no significant absorption above ~300 nm, showing that this protein purifies from *E. coli* without a heme cofactor, indicating a large drop in heme affinity. Signal is normalised to facilitate comparison. (*C*) UV-visible spectra of hemophilin apo proteins prepared by removal of heme in acid acetone and refolding (dashed lines): apo hemophilin (3.1 μM; black), apo hemophilin(H89Q) (3.0 μM; red) and apo hemophilin(H119Q) (2.3 μM; blue). Spectra are also shown after addition of ferric heme (2.3 μM; solid lines). The similarity of holo hemophilin and hemophilin(H89Q) spectra at all wavelengths, including Soret absorption band at 413 nm and α and β absorption bands at 538.5 and 574.5 nm, suggests that both proteins bind heme as low-spin Fe(III) species. Mixing heme with the H119Q mutant gave a spectrum with peaks at 404, 485 and 602 nm (Fig. S14), indicative of high-spin heme, similar to spectra of Fe(III) heme:protein complexes without an Fe-coordinating side chain (51) and monomeric heme in methanol:water mixtures (52). All spectra recorded in 0.2 M Tris.HCl, pH 8.0. (*D*) Agar well diffusion assays of apo hemophilin proteins in duplicate. Protein load is 20 μL at concentrations of: 1.25 μM (a), 2.5 μM (b), 5 μM (c) and 10 μM (d).

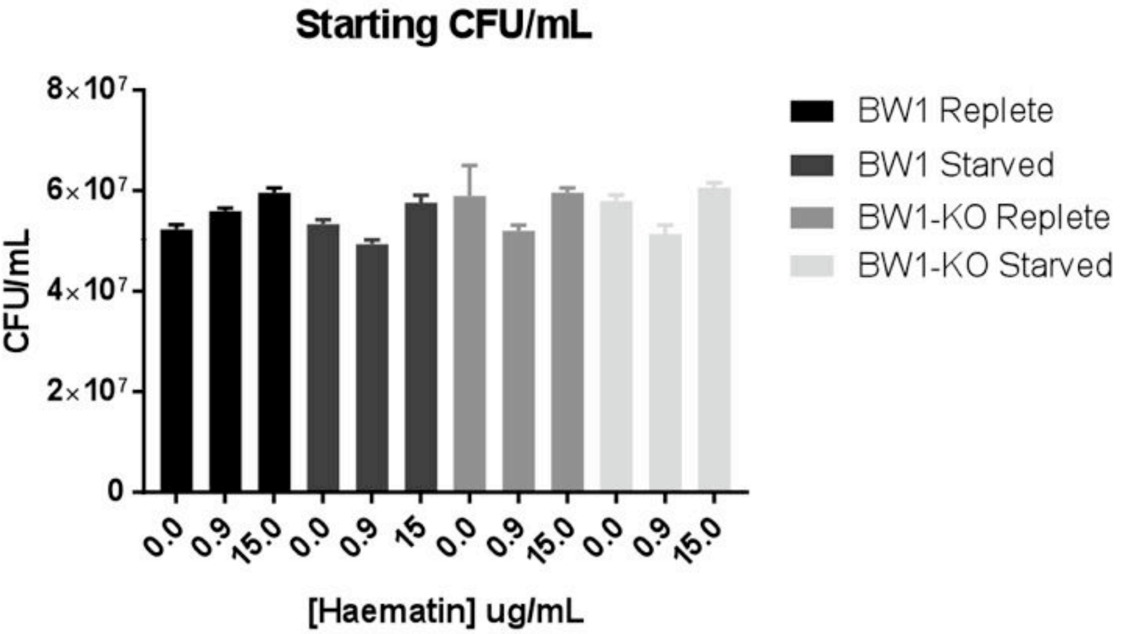


**Supplementary Fig. 14. Viability of cultures seeded in Figure 5.** Colony counts were performed on all suspensions by plating on CA to demonstrate that the pre-incubation period did not reduce cell viability and that equal numbers of cells were inoculated into the test conditions. Error bars represent ± SEM (n=3).

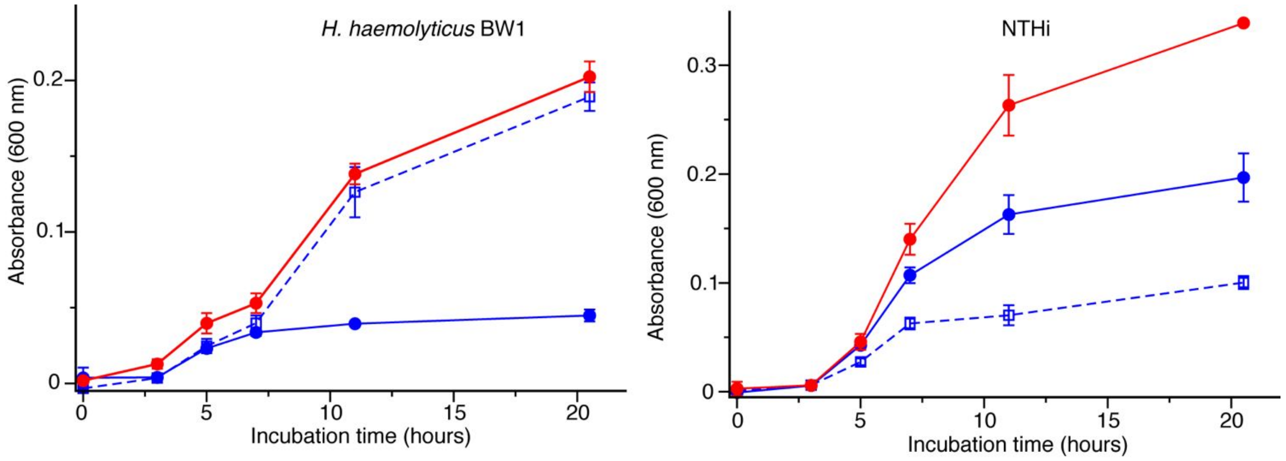


**Supplementary Fig. 15. The effect of heme-loaded hemophilin on the growth of Haemophilus species.** Growth on TSB supplemented with heme at 0.6 µg/mL (blue, solid line). Growth on TSB supplemented with heme at 5.6 µg/mL (red, solid line). Growth on TSB supplemented with heme at 0.6 µg/mL plus holo hemophilin at 7.7 µM (equivalent to 5 µg/mL heme), giving a total heme concentration of 5.6 µg/mL (blue dashed line). Error bars are mean ± SD (n=3).

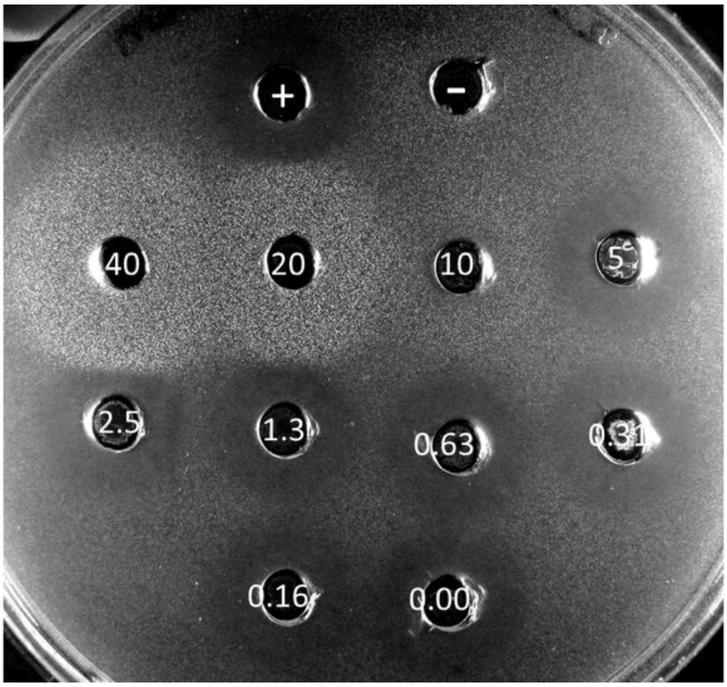


**Supplementary Fig. 16. Addition of heme neutralises the inhibitory activity of hemophilin.** Agar well diffusion assay with wells containing hemophilin from strain BW1 mixed with heme at variable concentrations (values shown, µg/mL). Positive control (+) contains BW1 hemophilin only. Negative control (–) contains DPBS. Indicator is NTHi strain NCTC 11315.

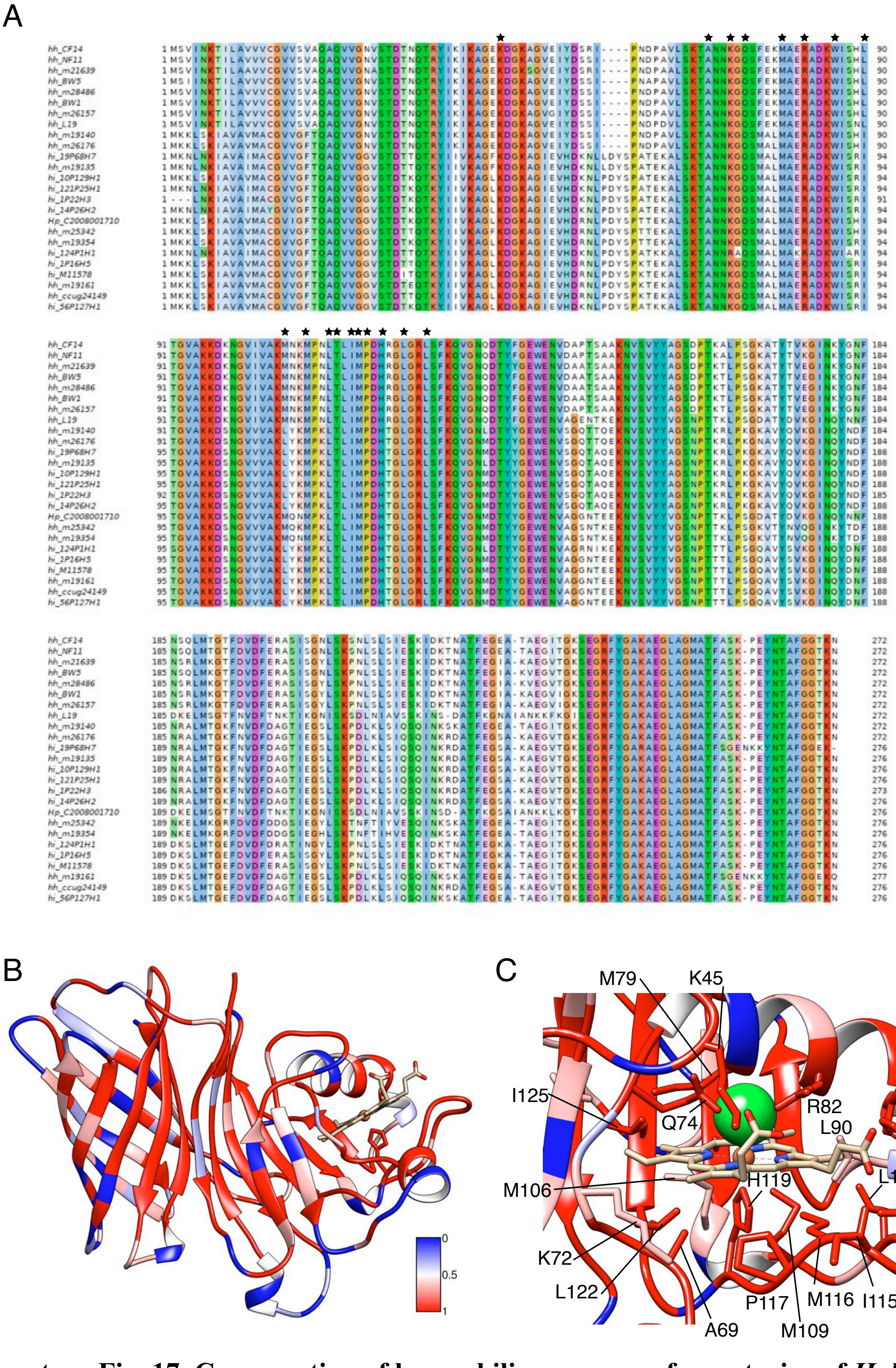


**Supplementary Fig. 17. Conservation of hemophilin sequences from strains of *H. haemolyticus*.** (*A*) Multiple sequence alignment of 25 unique hemophilin protein variants derived from strains of *H. haemolyticus* (strain names prefixed *hh_*), *H. influenzae* (prefix *hi_*), and *H. parainfluenzae* (prefix *Hp_*) that represent the full amino acid sequence diversity in all strains with a hemophilin gene at the time of writing. Star symbols indicate residues that make contacts with the heme group. (*B*) Ribbon structure of hemophilin coloured to represent amino acid conservation by physicochemical properties on scale ranging from 0 (dark blue, no conservation) to 1 (dark red, identical). (*C*) Conservation of residues in contact with the heme group, coloured as in B.

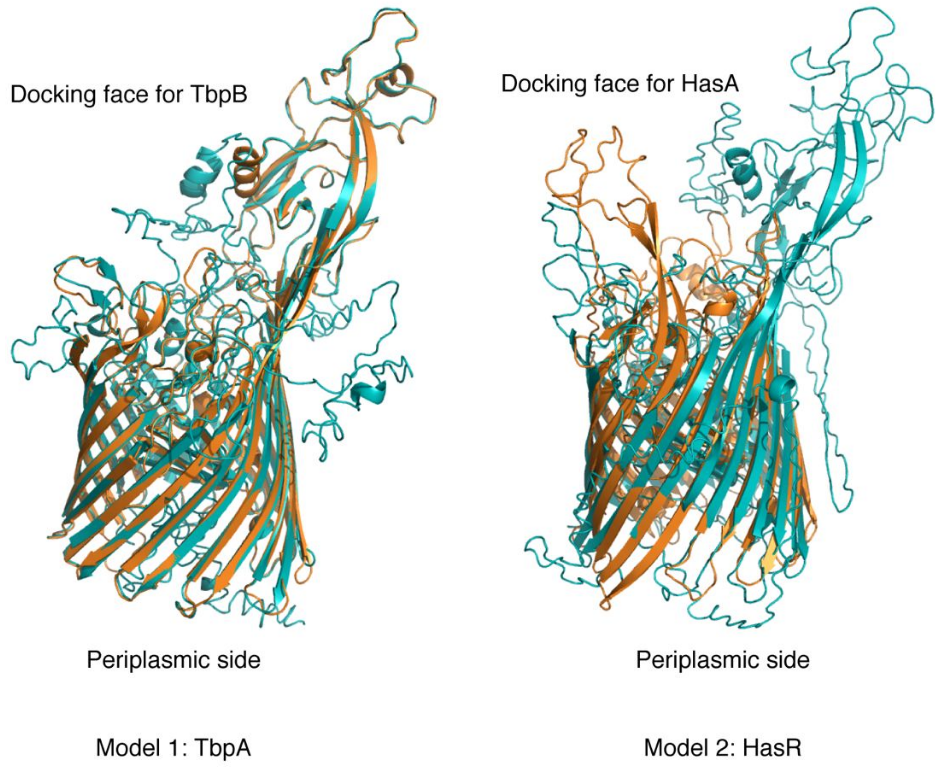


**Supplementary Fig. 18. Models of the putative hemophilin receptor.** Structural models of a predicted TonB-dependent transporter ORF immediately upstream of the hemophilin gene in *H. haemolyticus* strain M28486 (see Supplementary Table S3) were prepared using the PHYRE and I-TASSER web servers. This query sequence was chosen for modelling because the hemophilin gene from M28486 is 100% identical to that in BW1/RHH122. The top ranking template structures identified by both servers were the transferrin binding protein A (TbpA), and the receptor for HasA hemophore (HasR). The top-ranking model (Model 1) from PHYRE (orange) aligned 73% of the query sequence with TbpA (pdb 3v89 (53); 15% sequence identity over 811 residues) with a confidence score of 100%. Model 1 from I-TASSER used the same template but built more residues into the final model (cyan) with a C-score of –2.11 (acceptable range –5 to 2, which higher score indicating higher confidence) and TM score of 0.45±0.15 (TM>0.5 indicates shared topology). Model 2 was based on the HasR receptor using template pdb structures 3csl (PHYRE; 22% sequence identity over 721 aligned residues; orange) or 3ddr (I-TASSER; cyan) (54). Overall, Models 1 and 2 are similar with the greatest confidence in alignment in the transmembrane 22-strand β-barrel structure.

It is tempting to speculate that a conserved mechanism of interaction between hemophilin/HpuA/TbpB and their respective TonB dependent receptors, coupled with divergence of hemophilin/HpuA/TbpB ligand binding, might explain how these proteins have evolved to transport heme or iron from different host protein sources to the bacterial periplasm. Once delivered to the periplasm, there are several substrate binding proteins (SBPs) in *Haemophilus spp.* that can bind promiscuously to heme, peptides, glutathione, and other substrates (55-57), targeting these cargoes to ABC transporters that shuttle them across the plasma membrane.
